# Evolution of visual time throughout goal driven action learning

**DOI:** 10.64898/2026.01.22.701104

**Authors:** Nicola Binetti, Federico Mancinelli, Marco Zanon, Domenica Bueti

**Author notes:** dual senior authorship (Nicola Binetti, Domenica Bueti).

## Abstract

Timing is intimately woven into human cognition and behaviour, underscoring people’s ability of comprehending speech, playing instruments and competing in sports. Converging evidence suggests that our perceptual experience of time is grounded in our ability to control actions: time processing leverages motor control neural machinery, and perceived time is significantly distorted in proximity of action. However, the dynamic interplay between timing and action systems remains unexplored, particularly in goal-directed contexts where action control is gradually refined through practice. Through combined behavioural, neural and modelling approaches, we investigate how time processing gradually evolves as participants gain experience in goal-driven manual reaching actions, varying in terms of action-plan specificity. Participants judged brief visual durations while preparing hand movements aimed at visual targets displayed on a monitor. Subjective time expanded in the foreperiod preceding action onset, and this effect increased over trials, mirroring gradual improvements in action readiness as motor plans became more rapidly and efficiently deployed. Representation Similarity Analysis, integrating electroencephalographic and behavioural data, reveals that actions shape timing through independent mechanisms: planning specific behaviours shapes how duration information is written into timekeeping neural circuits as stimuli unfold (encoding). Parallely, more general aspects of motor engagement and anticipation shape how duration information is read-out from timing circuits following stimulus presentation (decoding). These findings highlight a dynamic, learning-driven coupling between perceptual time and motor control, where subjective time is recalibrated as goal-directed action preparation is optimized, shaped through independent time encoding and decoding mechanisms.

**Significance statement:** Despite lacking dedicated sensory and neural systems for duration, humans reveal a highly developed ‘sense’ of time, for instance when noticing subtle tempo changes in music. We show that perceptual understanding of time is linked to the ability of learning and fine-tuning action: our internal metric of time shifts as action preparation processes are gradually refined, affecting how quickly behaviours are planned and executed. This interaction is mediated by independent neural processes affecting the encoding (online monitoring) and decoding (offline readout) of duration. These findings shift our understanding of timing, from a passive sensory readout to one that is actively constructed and finetuned through action learning, and provide a mechanistic account of neural processes linking action preparation to time processing

## Main text

Timing is pivotal to cognition and behaviour, allowing people to predict events based on temporal expectations, to infer causality based on proximity of events in time, and to successfully interact with the environment. Humans exhibit highly precise time estimation, especially in millisecond to second ranges (1, 2), despite lacking dedicated sensory and neural systems for time. Converging evidence suggests that the brain bypasses these constraints by exploiting motor control circuitry, endowed with millisecond-level temporal precision required for action control (3). Indeed, at its core, action control can be viewed as a temporal problem, given that actions ranging from walking to striking a match-winning penalty kick, hinge upon precisely scheduled muscle activations.

A motoric basis of time is supported by neural and behavioural evidence. Brain studies highlight a spatially distributed timing architecture, rooted in a motoric ‘core timing network’ (4, 5), comprising the Supplementary Motor Area (SMA) (6), the basal ganglia (BG) and cerebellum (7, 8). This core set is augmented by auxiliary cortical and subcortical areas that are flexibly recruited based on task demands (9). Timing computations are similarly distributed in the frequency domain, carried by oscillatory neural activity involved in a breadth of motor related computations, such as action scheduling (10) (4-7Hz theta band), sensorimotor integration (11) (theta and 8-12 Hz alpha/mu) and prediction (12–14) (13-30Hz beta). A large body of behavioural studies also reveals a significant overlap between timing and action, where perceptual time is significantly distorted in proximity of behaviour. Action-timing interactions can be parsed based on when timing is sampled relative to action. Time is expanded as we prepare actions (11, 15, 16), whereas it is compressed while we carry out responses (17–19), and again expanded in the ensuing period where we evaluate behavioural consequences (20, 21). These effects interact with the way actions are carried out, modulated by direction (22), force (23, 24), frequency (25) and speed (26), generally reflecting a compatibility of action and perceptual magnitudes, with stronger, faster and larger movements expanding perceived time (27, 28). Studies also reveal that actions affect perceptual timing precision, with timing estimates sharpened during the execution of voluntary (29) and involuntary (30) behaviours.

While highlighting a complex and multifaceted relationship, timing-action interactions have mostly been explored through paradigms that overlook critical aspects of motor function, namely that actions are frequently goal-driven, and action control is gradually refined through practice. Actions involve goals (such as grabbing a set of keys laying on a table), plans that jointly specify motor instructions and predicted sensory consequences of behaviour (planning a reach and what we expect to feel when touching the keys) (31), and error signals (accuracy), critical for monitoring and course-correcting behaviour (32). Error signals are additionally instrumental to action learning, improving performance through gradual refinement of motor plans and control strategies (33, 34). However, most action-timing studies lack precisely defined action goals with associated errors, with participants performing behaviours with little or minimal constraint on movement parameters, such as moving in a given direction (22, 35), moving fast (26), freely exploring an action space (36), or involving simple actions operating at ceiling-level performance such as touching a screen (Hagura et al., 2012), or saccading to a visual target (37)). Tasks lacking explicit goals, or providing limited headroom for improvement, do not allow tracking changes in action performance across time, and measuring impact of these changes on perceptual time. Also, most action-timing studies provide static snapshots of timing performance, in terms of average timing estimates observed in an experimental condition relative to controls. However action control is refined over time, and given a motoric foundation of timing, one should expect variations in time perception throughout goal-driven motor tasks. Recent studies provide a significant stride forward showing that rhythmic motoric structure spontaneously emerges as individuals gradually refine skills at a throwing task (38), suggesting that implicit timing is an emergent property of goal-driven motor learning, and that implicit timing rehearsed in a motor task generalises to perceptual contexts (39).

Through combined behavioural, brain recording (Electroncephalography - EEG) and modelling approaches, we aimed to study how conscious (explicit) timing estimates are progressively shaped throughout goal-directed action control. We investigated how visual time is affected during action preparation stages in a dual hand-reaching / time-estimation task, with explicit action goals and associated error signals, and assess how this relationship evolves throughout trials as a function of changes in action performance.

## Results

An acoustic Go signal prompted participants to perform a ballistic hand reach aimed at circular targets displayed on a monitor placed horizontally above their dominant hand. On-screen target locations varied based on direction (Directional task: left or right), or distance (Distance task: rightward near or far), relative to fixation. Reaching behaviours required rapid horizontal shifts in index position, and were performed and tracked on a plexiglas surface placed under the monitor (Figure 1a-b). A visual depiction of the index landing point was displayed on the monitor at the end of each trial, providing accuracy feedback. Targets were either shown (Cued location amongst two alternative locations) or not shown (Non-cued location amongst two alternatives) during the action foreperiod preceding the Go signal (Figure 1d). Thus, Cued trials allowed for preparing a specific reach prior to the Go signal, whereas Non-cued trials, where target presentation coincided with the Go signal, additionally required participants to select amongst plans required in the current task (left or right in Directional; near or far in Distance task). Participants timed sub-second visual probe stimuli (.64∼1.24s) presented during the action foreperiod. The timing task was also performed without motor responses in matched intervals, requiring delayed timing judgements following the Go signal (Sensory control).

**Figure 1:**
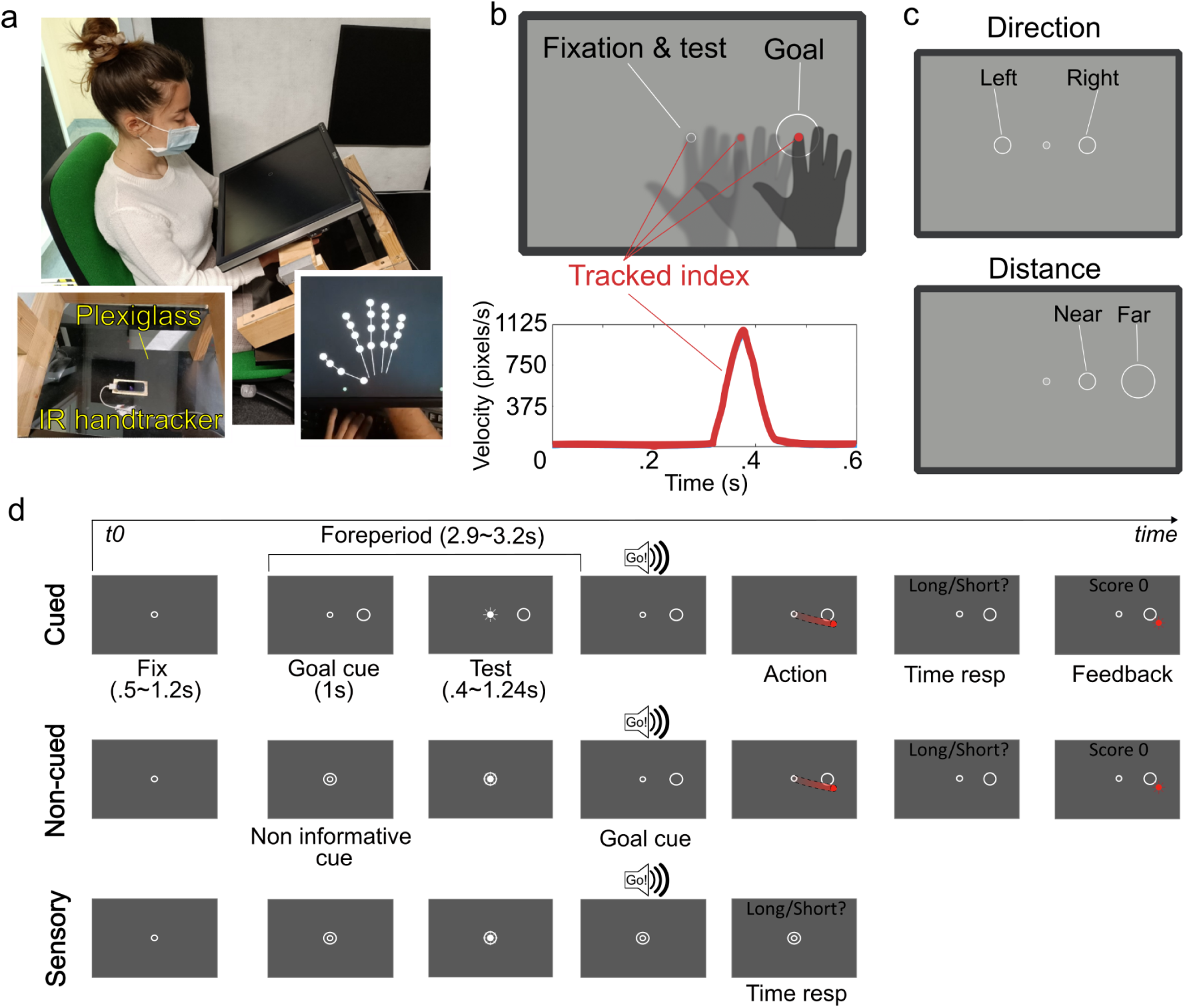
Task setup and experimental conditions. a) Participants performed ballistic horizontal hand movements on a plexiglass surface, tracked by an infrared (IR) camera. b) Movements were aimed at Goal locations displayed on a monitor that occluded participants’ hands, in Cued and Non-cued task conditions. Participants’ index remained stationary under the fixation point during matched passive observation conditions (Sensory). c) On-screen target locations varied based on direction (Directional task: left or right), or distance (Distance task: rightward near or far). d) During a variable foreperiod, participants were shown a Goal location in Cued trials (right or left in the Direction task; right near or far in the Distance task), and the fixation point lit up for a variable amount of time, providing a test duration. An acoustic beep (Go!), coinciding with the presentation of the Goal in the Non-cued condition, signaled the end of the foreperiod, requiring participants to sequentially perform the hand movement (Cued & Non-cued - index position occluded) and the timing judgement (all conditions). The final position of the index was revealed at the end of the trial, providing feedback on reach accuracy. A score (1-100) was provided for actions that landed within the goal area, proportionally to distance from the goal centre.

Visual time was perceptually expanded by action planning. On average, timing estimates expanded prior to action onset in Cued compared to Non-cued actions, and Sensory conditions. However, trial-wise models of timing responses revealed that both action conditions (Cued and Non-cued) showed progressive expansion of perceptual time across trials as action latency (the time required to initiate a reach following the Go signal, reflecting readiness to enact a motor plan). Representation Similarity Analysis (RSA) of EEG data showed that these distortions occurred during neural encoding and decoding of duration information, shaped by action planning and more general action preparation processes, respectively.

### Average Timing performance - action planning biases perceptual time

We compared measures of timing bias (Point of Subjective Equality - PSE: Test duration that is perceptually matched to the Standard) and sensitivity (Just Noticeable Difference - JND: detectable difference between Test durations) across Conditions (4 Cued conditions + Non-cued & Control), within both Directional and Distance tasks (Figure 2a). Linear Mixed Effects analysis on PSEs showed a perceptual expansion of time across Cued conditions relative to both Non-cued trials and sensory Controls, in both Directional (F(5,95)= 6.86, p=1.65e^-05^ ; R^2^=.06, conditional R^2^=.79; Figure S1) and Distance tasks (F(5,95)=3.15, p= 0.01; R^2^=.04, conditional R^2^=.7; Figure S2). Therefore when averaging across trials, Cued conditions, where motor plans were ready to be executed at the GO signal, were associated with a perceptual expansion of time, relative to Non-cued reaches that required plan selection at the Go signal, or matched controls that did not engage the motor system. Action features (left/right or near/far reaches), and hand-to-screen space mappings (congruent or incongruent mappings: see Methods) had no modulatory effect on timing estimates.

**Figure 2:**
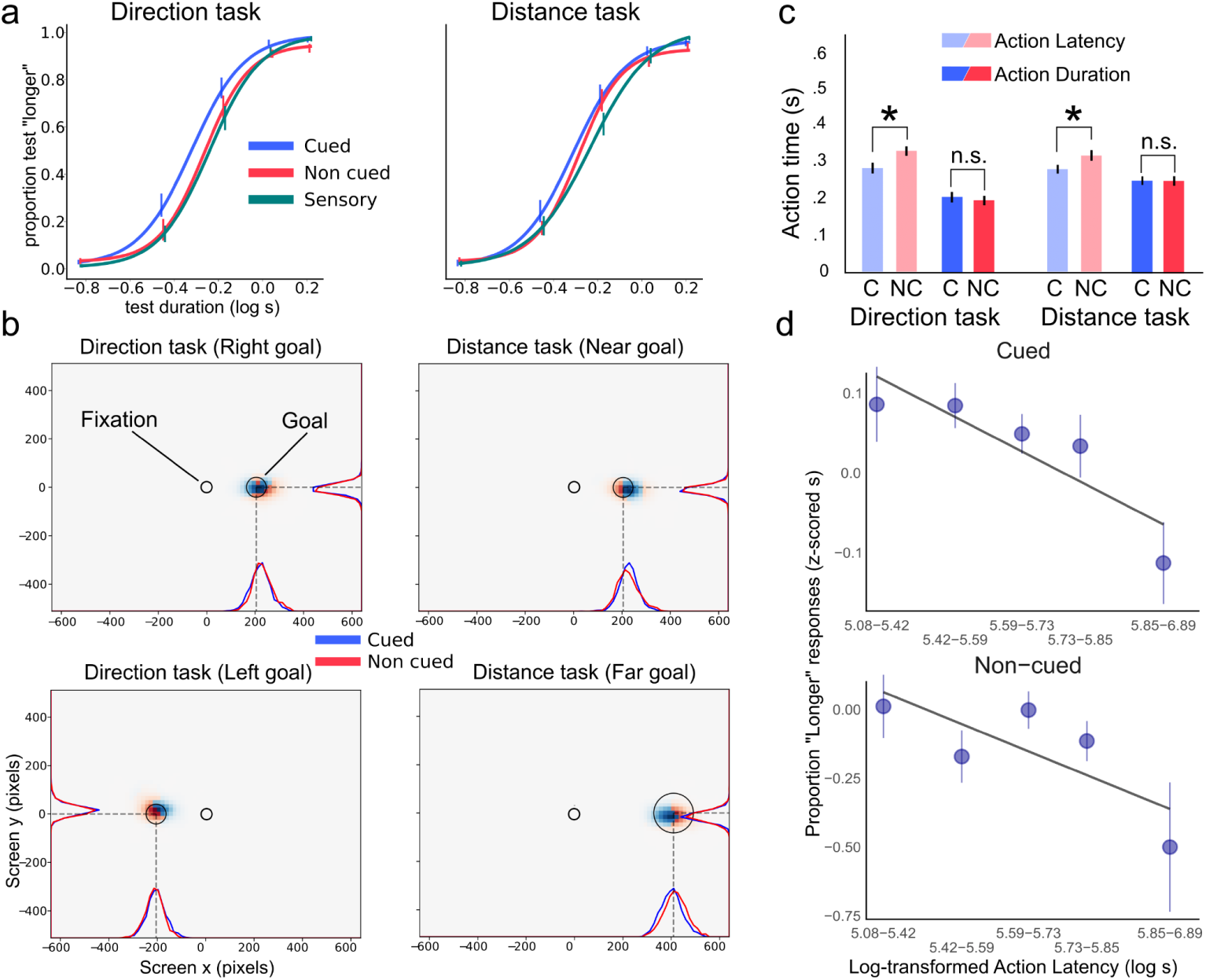
a) Psychometric fits of test “longer” responses as a function of test duration, across experimental conditions. Time is perceptually expanded in Cued relative to Non-cued and Sensory conditions, across both Direction and Distance tasks. b) Density of index finger landing points in screen space across experimental conditions and tasks (also projected onto individual axes). Action precision, quantified as distance between landing points and goal centre (which also inform reinforcement schedule) did not differ between Cued and Non-cued actions and between tasks, when accounting for linearly scaled errors for far goals in the Distance task. c) Action latency (time required to initiate action, from Go signal to movement onset) and duration (time required to complete action, from action onset to offset). Action latency was significantly lengthened in Non-cued (NC) relative to Cued action (C) actions due to goal location uncertainty (See Figure S3 and Figure S4). d) Perceived time (proportion ‘longer’) as a function of action latency. Shorter latencies were associated with longer duration percepts.

JNDs showed no significant difference in sensitivity to duration across conditions, in both Directional (F(5,95)=.43, p=.82; R^2^=.01, conditional R^2^=.24), and Distance tasks (F(5,95)=2.18, p=.06; R^2^=.06, conditional R^2^=.3). Thus timing task difficulty did not significantly differ across tasks (Directional or Distance) and conditions.

### Changes in time perception driven by action readiness

We modelled timing judgements on a trial-by-trial basis in Cued and Non-cued conditions to capture the dynamic relationship between changes in action performance and changes in time perception. We extracted action features from right index position time series, such as action latency, mean velocity, reaching error (the difference between the index’s final position and the goal center, reflecting reach precision), and distance traced (product of action duration and mean velocity). Individual factors for each test duration were inserted into the model so that we could assess the contribution of action features in explaining the residual variance.

Analysis revealed a strong influence of log-transformed action latencies on timing judgements (b = −0.79, CI_95 [−0.97, −0.62]). Shorter latencies were associated with increases in “probe longer” responses (time expansion) (Fig. 2c,d). Additionally, we observed significant effects of both task (Direction and Distance) and condition (Cued and Non-cued) on time responses. The Distance task resulted in greater time contraction (b = −0.20, CI_95 [−0.29, −0.11], relative to Direction), and Non-Cued condition resulted in greater time contraction (b = −0.30, CI_95 [−0.41, −0.19], relative to Cued). Independent of task and cue condition, log-transformed action latencies consistently decreased across trials (b = −0.05, CI95% [−0.06, −0.03]), while “test longer” responses increased correspondingly (b = 0.29, CI95% [0.12, 0.44]). Given these opposite changes, we next explored whether the rate of decrease in action latencies could predict the increase in perceived time expansion at the participant level. To address this, we constructed simple linear models of the trial-by-trial trajectories for both perceived duration and action latencies, incorporating participant-level random effects. To enhance statistical power for this individual-level analysis, data from the two tasks and cue conditions were combined, providing a single, robust random effect estimate per subject.

Participants’ individual parameters for rate of change (alpha parameters) in timing judgements across trials negatively correlated with rates of change in action response log-latencies (r = −0.32, p = 0.02, CI_95 [−1, −0.057]; Figure 3a). Participants with greater rates of decrease in action latency exhibited greater increases in ‘probe longer’ responses across trials (Figure 3b, c). As a sanity check, to reduce the impact of changes in action latency occurring early in each condition, we additionally excluded data from the first block in both conditions, focusing only on later trials. We then re-fit the model, generating new rate of change parameter estimates. This preprocessing step yielded consistent results, suggesting that the observed relationship reflects a task-wide process rather than transient performance fluctuations occurring in early trials only (r = −0.31, p = 0.03, CI_95 [−1, −0.04]).

**Figure 3:**
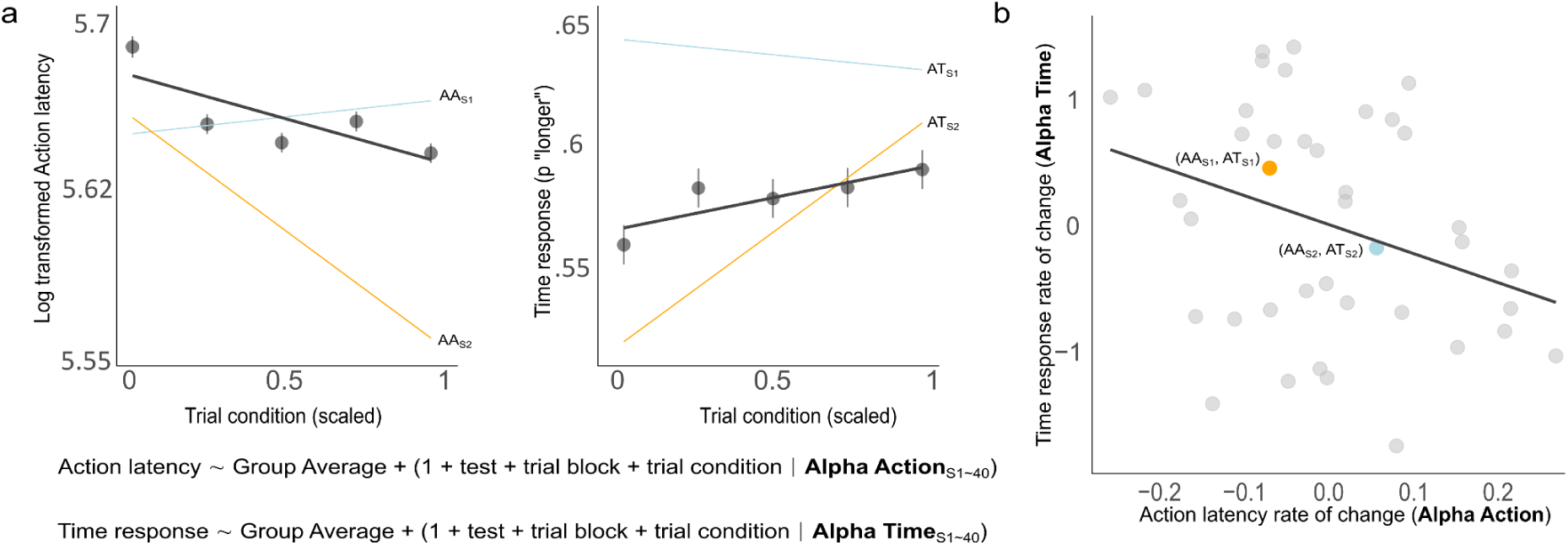
a) Group Average changes (thick dark line) in Action latency and Time responses (“longer” judgements) as a function of trial number throughout Cued and Non-cued trials (scaled to account for differences in trial numbers). Action latency and Time responses were separately modelled as a function of trial number within each condition (capturing global changes occurring across all Cued/Non-cued condition trials - significant) and trial number within blocks of 20 trials (capturing local changes associated with switches between task conditions - non-significant). Participants were modelled as random effects (coloured lines), providing individual parameters for rate of change in action latency (Alpha Action) and time responses (Alpha Time) across trials, controlling for other sources of variation. b) Alpha Action and Alpha Time parameters across participants, showing inversely related rates of change in action latency and time responses.

### EEG Time Encoding and Decoding Correlates

ERPs for Time Encoding and Decoding were analysed to identify the effect of Test Duration (5 probe durations) and Condition (Sensory, Non-cued and Cued) in both groups separately. No significant effects were observed during Time Encoding, for Test Duration nor for Condition (Figure 4a, left column). Since any possible information about probe duration is provided during the selected interval for ERPs (in fact, epoches ranged from the probe onset to the offset of the shortest probe), no effects for Test Duration were expected. Whereas, lack of differences among conditions suggest that action planning does not seem to set a condition-specific “state” for time encoding.

**Figure 4:**
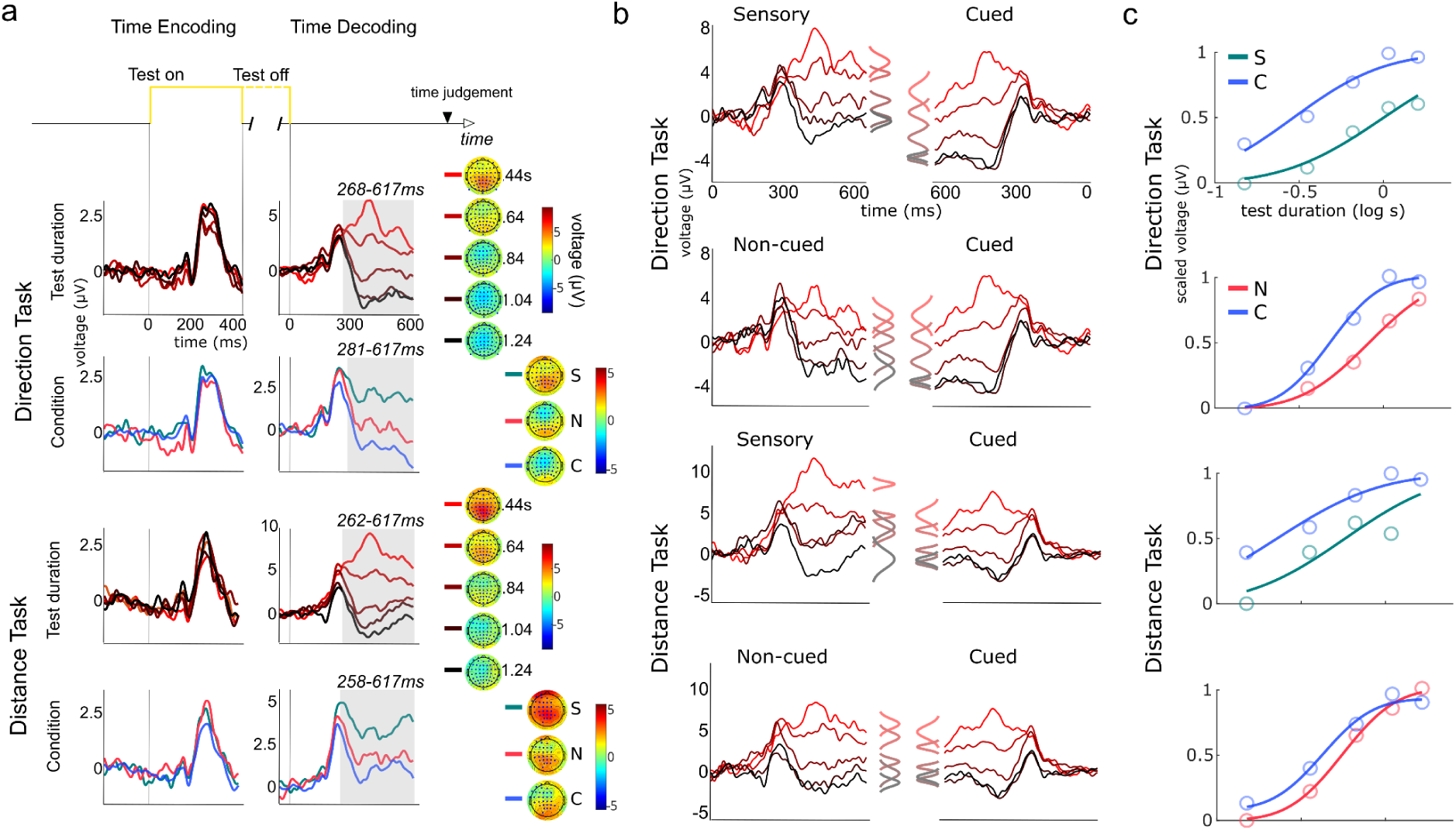
a) Evoked neural responses time locked to test duration onset and offset, indexing time encoding (online monitoring) and decoding (offline readout), respectively. The time encoding interval was constrained to the shortest tested duration (420ms), while the decoding window corresponded to the shortest interval afforded by the variable foreperiod preceding the Go signal (617ms). Gray shaded areas show intervals with significant differences between test durations (.44-1.24s, averaging across conditions) or conditions ((S)ensory/(N)on-cued/(C)ued, averaging across test durations), with corresponding topography of evoked signals on the right. b) Side-by-side comparison of time decoding in Cued Vs Non-cued/Sensory conditions (flipped on the time axis) at electrode Cz. Gaussian curves represent the average voltage +/- scaled variability within time windows showing significant differences between tested durations (shaded regions in a), revealing a reduction of voltage across tested durations in Cued relative to Non-cued & Sensory conditions. c) Neurometric curves (cumulative Gaussian fit with lapse rates) of average normalized voltage within significant time decoding windows (shaded regions in a) as a function of tested duration, for (S)ensory, (N)on-cued and (C)ued conditions.

Significant effects were observed during time Decoding, in both groups. The effect is significant about 270 ms after probe offset, with a peak at about 350 ms (Figure 4a, right column), in central and parietal electrodes. ERP amplitudes depend on test duration: the shortest duration (0.44 s) elicits a large positive potential, which diminishes with longer durations, showing an exponential decay (Figures S7 and S8). The result is in line with previous findings. Ofir and Landau (2022) showed that the EEG signal recorded in fronto-central electrodes about 300 ms after probe offset is a neural marker of the readout of the time perception system that during duration encoding accumulates evidence in favor of a “longer” response. To note, clusters of significant time points and channels were observed also for earlier latencies, but these are likely byproducts of the baseline correction. In fact, the 100-ms baseline interval for Decoding epochs was centered on probe offset, an interval that still includes ERP components elicited by the onset of the stimulus (See Figure S9; also present in Ofir and Landau 2022).

The time Decoding effect of Condition is observed at frontocentral electrodes, where Non-cued and Cued conditions elicited more negative potentials than Sensory in both groups. These differences emerged around 300 ms following test offset. The similar ERP amplitudes in Non-cued and Cued conditions, both distinct from Sensory, suggest that action readiness processes drive ERP components from this time point onward. This interval appears crucial for understanding the neural basis of the behavioral bias in time perception. However, it also marks the overlap of significant effects of both Probe Duration and Condition. At this stage, participants are preparing to move, especially in the Cued condition, where early cue presentation allows planning upcoming behaviour. Consequently, ERP activity related to motor preparation may obscure subtle responses linked to time perception, which is behaviorally biased by motor preparation activity. To disentangle these overlapping processes, Representational Similarity Analysis (RSA) was employed. Still, ERP comparisons across the three Conditions support the idea that Motor conditions (Cued and Non-cued) are well matched.

### Neural source of distorted percepts of duration

We aimed at identifying the neural source of perceptual distortions of time using Representational Similarity Analysis (RSA), identifying brain correlates based on similarly patterned behavioural and EEG activity (Figure 5).

**Figure 5:**
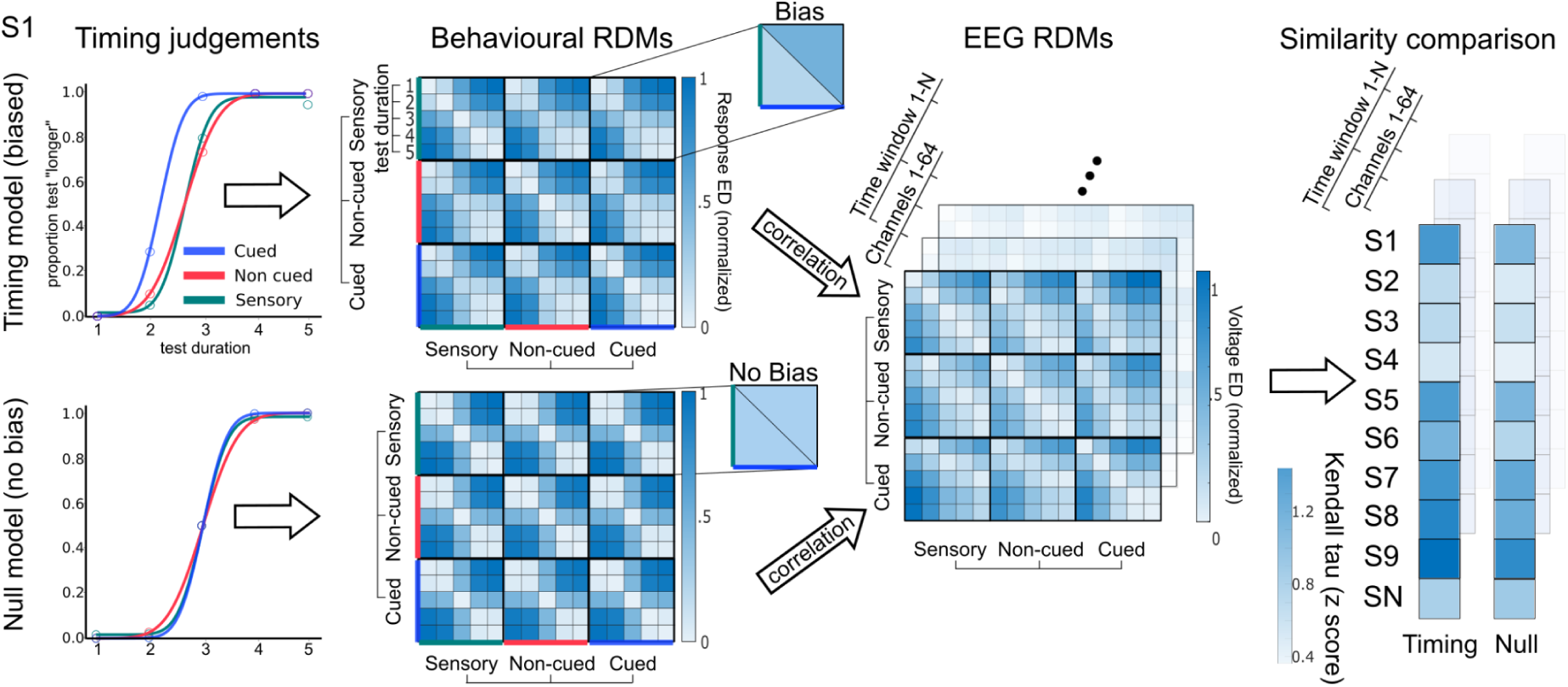
Representation Similarity Analysis pipeline used to reveal the neural source of the time perception bias observed in Cued relative to Non-cued and Sensory conditions. Behavioural RDMs were built based on individual participant data (Euclidean Distance in participant time judgements) or synthetic data (Euclidean Distance in unbiased participant time responses - providing a null model), capturing the overall behavioural performance across tested durations and experimental conditions. Participant Behavioural RDMs were compared to EEG RDMs (Euclidean Distance in voltage, per channel and 50ms time window) via Kendall Tau correlations, indexing pattern similarity (z scored) between real/synthetic behaviour and neural response. Real and synthetic pattern similarity z-scores were compared via t-tests, revealing channel/s and time window/s at which behavioural/neural similarity was greater for biased relative to unbiased time perception, i.e. identifying where and when neural response more closely resembles biased time perception, relative to unbiased perception.

On average, perceived time was significantly lengthened in Cued relative to Non-cued actions and Sensory controls, for both Direction and Distance tasks. RSA results reveal that the perceptual expansion of time was associated with biased neural activity in time encoding and decoding, where duration information is respectively monitored and read out for perceptual judgements. Time encoding and decoding responded differently across experimental conditions (Figure 6; see also Figure S10-S13). In the Direction task, time encoding was specifically impacted by the presence of a motor plan, as captured by contrasting Cued and Non-cued actions, reflected in responses at central (287-336ms) and bilateral posterior electrodes (383-432ms), following probe onset. In the Distance task, we also observed effects on time encoding: biased perception in Cued respect to Non-cued actions, was reflected in responses at central (50-100ms) and temporo-parietal electrodes (336-383ms), following probe onset. In the Direction task time decoding was more generically linked to motor system engagement, captured both by Cued Vs Non-cued and Cued Vs Sensory contrasts, and reflected in responses at frontocentral (359-412ms) and central electrodes (308-463 ms) respectively, following probe offset. The Distance task also showed similar patterns of activity in time decoding, which however did not reach significance.

**Figure 6:**
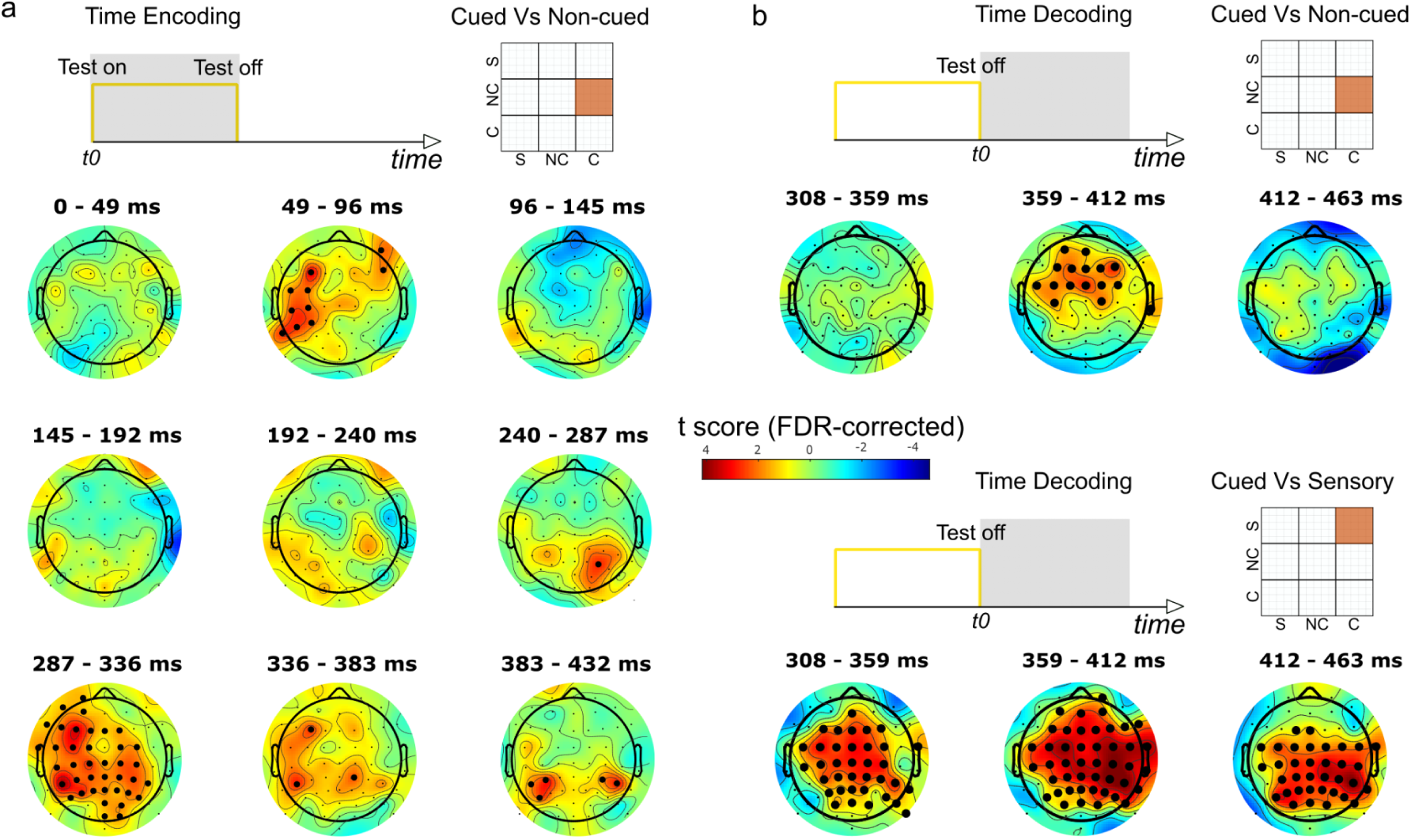
RSA contrasts targeting RDM sub-quadrants, identifying neural correlates of perceptual distortions of time within time encoding and decoding windows. Perceptual biases in timing were reflected in neural activity throughout time encoding and decoding. a) RSA revealed effects on time encoding specifically linked to action planning (Cued Vs Non-cued; summarizing effects in both Direction and Distance tasks, individually modelled in SI), within windows time-locked to Test onset. b) Effects on time decoding more generically linked to motor system engagement (Cued Vs Sensory and Cued Vs Non-cued, only significant in Direction task), within windows time-locked to Test offset.

## Discussion

Action and timing systems are deeply integrated in the brain. From a timing perspective, our ability of estimating duration engages motor systems (40, 41), even in tasks requiring no overt behaviour (5, 28, 42), and time perception is profoundly shaped by processes that straddle the line between action preparation, online control and post-hoc evaluation of action consequences (18, 22, 24, 43). Action control in itself is a temporal (i.e. scheduling) problem (3, 44). Logically, this implies that time perception exploits systems that are pre-equipped for precise timing (45). Action and timing are also inherently linked due to their dynamic nature. Actions are gradually honed through practice, driven by external goals and reward signals, while timing involves the integration of continuously unfolding duration evidence. Our findings highlight the dynamic interplay between action and timing, driven by the optimization of action selection and planning processes, that impact how quickly voluntary actions are initiated.

### Action planning expands visual time

Comparison of average time perception biases (PSEs) across conditions showed time expansion during Cued trials compared to Non-cued and Sensory controls, thus suggesting that priming specific motor plans distorts subjective time. Previous studies have linked time expansion to preparation of hand reaches (15), button presses (11), or horizontal arm movements (22). Hagura and co-workers showed that time expansion depends on the mere intention to act, as it also occurs when planned actions are aborted following a no-go signal (15). Iwasaki and co-workers showed that time expansion during action preparation cannot be attributed to generic attention / arousal effects, as this phenomenon only occurs for filled intervals (16) where visual information is continuously delivered, opposed to empty intervals signaled by visual transients at the start and end of tested durations. Thus, action preparation modulates processing of concurrent sensory information, as well as temporal features embedded in the sensory signal, while attentional effects would impact timing irrespective of how duration information is delivered. PSE differences between Cued and Non-cued/Sensory add to this literature, highlighting that time distortions prior to action are linked to action plan specificity.

The interaction between motor and sensory systems conceptually aligns with the embodied cognition framework (46), where action possibilities (plans, motor repertoire and affordances) are integrated with perceptual mechanisms. A more mechanistic view frames this interaction in terms of sensorimotor interaction processes, where action preparation primes sensory circuits in anticipation of upcoming changes, contingent on action execution (47, 48). These interactions are instrumental for aligning action and perception systems, discounting latencies and noise in information processing, and ensuring that sensory information is optimally integrated into behavior. Distortions in perceptual time can be viewed as a byproduct of interaction mechanisms that temporally integrate motor and sensory signals (47) and/or enhance processing of upcoming sensory consequences of behaviour (15, 49).

Spatial features of action have been reported to alter time. Tomassini and co-workers tested timing in the brief moments preceding rightward or leftward hand/arm movements, or during isometric (stationary) contractions requiring matched force output. The authors observed that movement/force direction affected timing estimates, with relative time contraction and expansion for leftward and rightward directions, respectively (22). We observed no effect of action features (L Vs R or Near Vs Far reaches) nor action mappings (congruent / incongruent action-visual outcome mappings) on visual time. Thus, time perception was modulated by the temporal aspects of planned actions (when planned action is initiated) opposed to non-temporal action parameters (how action is carried out). These differences could possibly depend on paradigm design. One possibility is that we are dealing with different processes along the motor hierarchy, i.e. earlier action selection (50) opposed to later activity triggering motor responses (51). Tomassini & Morrone randomly sampled timing performance (150ms standard) across a 200ms window between the Go signal and action execution, targeting processes that immediately precede motor responses. We probed timing between the target presentation and the Go signal, over a significantly longer interval (900ms standard), focusing on earlier action selection/planning (Cued, and Non-cued, accounting for target location uncertainty - see next section) or more general motor anticipation processes (Non-cued) leading up to responses. Early / later processes can afford different opportunities in terms of interactions between time and spatial components of action (52, 53). Differences between early and late preparation processes might also account for differences in time modulations observed between Tomassini (22) and Iwasaki and co-workers (16). Iwasaki and co-workers’ study, methodologically closer to our setup (targeting longer intervals, prior to a go signal), did not reveal timing modulations with empty intervals, whereas Tomassini and co-workers’ observed effects with empty intervals, thus suggesting different mechanisms. Another possibility relates to differences in tested duration, with a 900 and 150 ms standard duration, respectively for our and Tomassini and co-workers’ study. The latter noticed non-monotonic modulations in direction-timing interactions, based on when timing is probed relative to action, possibly reflecting periodic fluctuations in sensory processing time-locked to action (47, 54). Given the frequency of these fluctuations (theta range, ∼200ms period), 900ms intervals might prove ineffectual at registering finer interactions between motion direction and time perception.

### Perceptual time is dynamically shaped by changes in action latency

While on average Cued actions biased timing estimates relative to Non-cued/controls, trial-based timing models revealed a more nuanced relationship linking changes in action latency and time perception. Within participants, visual time progressively expanded as action latency decreased (faster reaction to Go), across both Cued and Non-cued trials. Given predefined target locations, Cued responses showed a marginal reduction in action latency across trials. Reduction in latency was more pronounced in Non-cued trials, likely because these offered a larger margin for process optimisation as participants learned to cope with target location uncertainty. A possible strategy for dealing with target uncertainty in Non-cued trials can consist in simultaneously planning competing actions, e.g. preparing both L and R reaches in the Directional task, and then initiating the correct response following target presentation, coinciding with the Go signal. Alternatively, action planning and execution can be purely reactive, triggered by the Go signal. Either way, both strategies can be optimized through practice, leading to faster selection, planning and initiation of actions. Independent of differences between Cued and Non-cued trials, decreases in action latency were accompanied by similarly scaled increases of perceived duration, suggesting a general link between trial-dependent improvements in response speed and changes in visual time.

Both approaches, comparing PSEs between conditions (average responses across trials) and trial-based models relating action latency and timing response rates of change, capture the same general effect. The difference in PSEs between Cued and Non-cued, reflect the overall lower latencies in the Cued condition, where the goal is clearly signaled at the beginning of the trial, allowing immediate action selection and planning. This overall advantage of Cued trials is reflected in the difference in trial-wise proportion of “longer” response intercepts between conditions (Figure S5). Reductions in action latency can have several underlying causes. These include optimized motoric processes as a function of practice and proceduralization, such as reduction in time required to select, plan and trigger behaviours (55), as well as non-motoric processes, such as increase in attention (56), cortical arousal (57) and dopaminergic activity (58, 59). Although the task itself offered no extrinsic primary rewards to reinforce reductions in response latency, engagement itself in the task might bear a time-wise opportunity cost that motivates subjects to streamline planning and action selection (59, 60). Despite different potential causes for reductions in latency, RSA analyses, linking action preparation to time perception, narrow down effects to brief windows of time encoding and decoding activity. No effects were observed in neighbouring epochs lacking timing computations, involving goal stimulus processing, ruling out non-specific effects of attention or motivation, which would be expected to influence a broader range of cognitive operations beyond temporal processing (discussed below).

While changes in response latency influence time expansion, and these changes can reflect optimization of action selection / planning stages (Diedrichsen & Kornysheva, 2015; Shmuelof et al., 2012), changes in action precision did not affect timing, despite similar overall improvements across trials (evidence of motor learning, Figure S6). A possible explanation for this is additional noise associated with action execution, which occurs on top of action planning. With practice, participants may plan their behavior more efficiently (i.e., they develop a clearer understanding of the required action given the current goal and mapping constraints); however, selected plans do not necessarily yield precise actions.

### Action planning shapes neural encoding and decoding of time

Timing mechanisms are indexed by distinct neural activity patterns, reflecting transformation of sensory duration information into perceptual judgments. The accumulation of timing evidence has been linked to various time-dependent neural responses, including ramping, tuning and speed of neural trajectories(61–63). Timekeeping is distributed across sensory, cortical, and subcortical structures, with sensory cortices locally encoding and storing temporal information embedded within sensory channels (64). Duration information propagates to higher-order regions such as the SMA, cingulate cortex, insular cortex, basal ganglia, cerebellum, and thalamus (6, 65, 66), where it is represented in modality-independent formats that inform perceptual and decisional aspects of duration (42, 67).

Analysis of EEG data showed spreading out of test duration-evoked responses (Figure 4a, right column), time-locked to time stimulus offset. Comparison of ERPs across tested durations showed monotonic increases in signal amplitude between linearly spaced tested durations, reflecting differences in accumulated timing evidence. This effect mirrors recent EEG data demonstrating distinct neural signatures of visual time, peaking at frontocentral electrodes 300–500 ms following the offset stimulus presentation (68, 69). This evoked activity, modeled as a function of stimulus duration and participant responses, provides evidence of time decoding linked to perceptual decisions. While replicating these results, our findings show that time-decoding ERP amplitudes were modulated by action conditions (Cued/Non-cued Vs Sensory; Figure 4b,c), thus suggesting that neural activity associated with motor preparation biases concurrent time decoding. ERP data was further examined using Representational Similarity Analysis (RSA), identifying the neural signature of perceptual distortions of time observed during action planning (relative to unplanned actions and control; Figure 5/6). RSA revealed that perceptual distortions of time are reflected in neural response during time encoding, specifically linked to motor planning, as exclusively captured in the Cued Vs Non-cued action comparison. Additionally, perceptual time was reflected in neural response during time decoding, more generally linked to motor system engagement, as captured in both Cued Vs Non-cued and Cued Vs Sensory comparisons. This difference might suggest distinct neural mechanisms underlying condition-specific distortions in time perception: encoding processes are selectively sensitive to the presence of action plans (for both Direction and Distance), while decoding captures both planning-related dynamics and broader motor system engagement (only in Direction). RSA also showed that timing biases were specifically linked to neural activity in brief windows within time encoding / decoding epochs, whereas no differences were observed between conditions in prior epochs associated with processing of the Goal stimulus. These results specifically link perceptual distortions of time to timing-related activity, opposed to neural processing of non-temporal information.

Our findings align with broader evidence linking action-timing interactions with sensorimotor integration mechanisms. Functional connectivity studies demonstrate that when visual events are triggered by voluntary actions, perceptual and motor processes synchronize through oscillatory coupling across primary motor and sensory cortices (47). By extension, sensorimotor integration directly influences duration information carried along sensory channels, with temporal distortions explained by phase-locked sensorimotor coupling. Our results contribute to this literature by showing ERP shifts that reflect action-planning specificity, i.e. action planning (captured on average by the Cued condition), opposed to more generic motor preparation in reactive / unplanned actions (Non-cued), or disengagement of the motor system during passive observation (Sensory control) (Figure 4a). These average shifts in activity provide different neural backdrops on top of which timing computations are embedded. RSA analysis links ERP differences to perceptual judgments, showing that these shifts in neural activity related to action preparation affect different stages of timing computations.

## Conclusion

Our findings reveal that perceptual time is not static, but is dynamically recalibrated through goal-directed action. As participants performed hand-reaching actions with varying degrees of planning specificity, perceived duration progressively expanded, driven not by execution accuracy, but by gradual decreases in the timing of action selection and initiation. Trial-wise coupling between time expansion and reduced movement latency highlights the role of motor planning temporal dynamics in shaping our perceptual sense of time. Crucially, EEG/behavioural representational similarity analyses show that these distortions arise during both the online monitoring (encoding) and offline readout (decoding) of temporal information, implicating action planning and more general aspects of motor engagement and anticipation, respectively. Together, these results position action planning as an internal metric for time estimation and establish a learning-sensitive framework linking perceptual time to the evolving dynamics of goal-driven behaviour. This work advances a unified perspective on how cognitive and motor systems interact to enable our perceptual experience of time.

## Methods

### Participants

We tested 40 right-handed adults with normal or corrected-to-normal vision, in two motor-timing tasks with EEG (Directional task N=20, 14 Females, 23.6±2.2 years old; Distance task N=20, 14 Females, 24±2.9 years old). Participants were recruited through the SONA online recruitment platform, and were compensated 25€ for 2.5 hours, which included hand tracking calibration, EEG cap and electrode setup while training at the action-timing task, and experimental data collection punctuated by short breaks. Participants viewed and signed information and consent forms prior to testing. The experiment was approved by the SISSA ethics committee and conformed to the guidelines of the declaration of Helsinki.

### Sample size

Sample size was informed by sample size calculations run on pilot data (.8 target statistical power), consistent with numbers tested in equivalent studies in literature (11, 48, 70).

### Apparatus

Participants performed a reaching task with their right index finger on a plexiglass surface mounted on a custom desk, while simultaneously estimating durations of brief visual stimuli. Hand position was visually occluded by the LCD monitor (DELL 1905FP, 19”, 1280x1024 pixels @ 75Hz refresh rate) positioned about 10cm over the participant’s hand, supported by a custom wooden bracket. An infrared (IR) hand tracking camera (Leap Motion Controller - https://www.ultraleap.com/leap-motion-controller-whats-included/) captured hand movements at 75Hz (constrained to monitor refresh rate). EEG signals were recorded with an ActiveTwo EEG system (Biosemi, Amsterdam, The Netherlands) at a sampling rate of 2048 Hz with 24-bit resolution. The Biosemi amplifier applied no high-pass filter (DC-coupled) and a low-pass 5th-order sinc filter, with a −3 dB attenuation at approximately 1/5th of the selected sampling rate, resulting in an analog signal bandwidth of 417 Hz at the selected sampling frequency. The signals were acquired from 64 active electrodes mounted on an elastic cap positioned according to the standard 10/5 coordinate system. Two additional electrodes replaced the ground electrode in conventional systems: the Common Mode Sense active electrode (CMS) and the Driven Right Leg passive electrode (DRL). The signals were recorded in single-ended mode (i.e., not referenced). Four additional electrodes were used to monitor eye movements. Specifically, two electrodes were placed on the outer canthi of both eyes to record horizontal movements (hEOG-left and hEOG-right), whereas two electrodes placed beneath and above the left eye (vEOG-lower and vEOG-upper, respectively) were used to monitor vertical movements and blinks. Finally, signals from the left and right mastoids were also recorded to compute offline the average-mastoids reference. The magnitude of the offset of all electrodes was held below 20 mV. The experiment was performed in a dark and sound-proof testing booth.

### Procedure

Participants initially performed a calibration routine to align IR camera and screen spaces by touching points with the index finger of the dominant hand on a 3x3 grid drawn on the plexiglass surface while the monitor was temporarily removed. Once calibration was completed, the monitor was secured to the bracket over the plexiglass surface. This was then followed by a training phase that allowed participants to familiarize themselves with the setup and experimental tasks, while EEG cap and electrodes were mounted. The training broke down the main components of the dual task, first focusing on reaching movements, followed by the temporal estimation task, and finally combining both tasks. In the reaching task participants had to initially position their index finger on a starting point on the plexiglass surface by aligning a small circle symbolising their index on the monitor, to a small cross in the central region of the monitor (position was randomly jittered across trials to avoid reliance on the monitor bezel as a spatial frame of reference). After reaching this position, the cross changed into an empty circle representing a fixation point lasting a variable amount of time (.5∼1.2 s), after which a circular goal area appeared at a horizontally displaced location relative to fixation for 1 second. In the Directional task this could be at 205 pixels (approximately 5.5cm) to the right or left of the starting point, or in the Distance task at 205 or 410 pixels to the right of the starting point. After a variable foreperiod interval ranging between 2.9 and 3.2 seconds, an acoustic Go signal (brief auditory beep delivered through computer speakers), instructed participants to perform a ballistic reaching movement aimed at the target location with their index finger. No online feedback on index position was provided throughout the movement, whereas position feedback was provided at the end of the reach, displaying the index’s point of arrival relative to the target area. Stimuli were displayed in grey against a black background. A score between 0 and 100 was displayed at the end of each trial above the goal area based on proximity to goal, to keep participants engaged and strive for improvement (scores were greater than 0 if the index was within the goal area, reaching 100 when the index landed on the goal area’s centre). Participants were informed that at the end of the experiment they would be shown their total score (sum of trial scores), ranked based on performance relative to other tested participants. Participants were trained at the motor task until they could reliably perform ballistic reaches while waiting for the Go signal.

Participants were then trained at the timing task, without performing motor responses. Participants were first shown a fixation point (empty circle) that was intermittently filled (filled white circle), providing 15 repetitions (jittered onset) of a .84 second Standard duration that participants had to memorize. Then, participants were asked to compare Test durations against the memorized Standard, based on the method of single stimuli. One second following the goal presentation, the fixation point was filled (filled circle) for a brief amount of time ranging between .44 and 1.24 seconds, in 5 linearly spaced intervals, providing a Test duration. After the Go signal was presented, participants indicated with a left or right mouse button press (held with their left, non-reaching hand), whether the Test appeared shorter or longer than the Standard duration (left or right button, respectively). Finally participants were trained with combined motor and timing tasks, thus requiring them to monitor the Test stimulus as they prepared the reaching action, and to judge its duration after the completion of the action. Participants were trained until they reached 100% accuracy at the timing task with the shortest and longest test durations (.44 & 1.24 second Tests).

The testing phase involved 3 conditions (counterbalanced across participants), where time was estimated while: 1) planning hand movements aimed at cued goal locations (Cued condition - priming a specific plan initiated at the Go signal, as done in training), 2) during the foreperiod of reactive movements aimed at Non-cued goals (Non-cued condition - a non-informative goal is shown on top of the starting point, requiring a selection amongst alternative action plans at the Go signal), and 3) during matched intervals involving no reaching movements (Sensory). The Cued condition involved 20 blocks of 20 trials each. Each Cued block consisted in congruent or incongruent trials, where we manipulated camera to screen-space mappings: e.g. in the Directional task a leftward hand movement produces a leftward (congruent) or rightward (incongruent) movement on screen space, or in the Distance task a small hand movement produces a small (congruent) or large (incongruent) displacement in screen space. Congruent and Incongruent blocks were interleaved (i.e. alternated every 20 trials), with a text prompt informing participants on the current block type. The Block order (starting with Congruent and Incongruent) was counterbalanced across participants. Thus Cued condition involved four types of trial for Directional task (Left Congruent - LC; Right Congruent - RC; Left Incongruent - LI; Reft Incongruent- RI) and for the Distance task (Near Congruent - NC; Far Congruent - FC; Near Incongruent - NI; Far Incongruent - FI). These manipulations were aimed at studying how planned action features and their predicted sensory consequences (which can be disentangled based on Congruent and Incongruent mappings), affect timing estimates. Non-cued and Sensory conditions were alternated in 5 blocks of 20 trials each (i.e. 20 Non-cued, 20 Sensory, 20 Non-cued etc). Cued Vs Non-cued/Sensory condition order was counterbalanced across participants.

### EEG recording

- EEG signals were recorded with an ActiveTwo EEG system (Biosemi, Amsterdam, The Netherlands) at a sampling rate of 2048 Hz and a 24-bit resolution. The Biosemi amplifier applied no high-pass filter (DC-coupled) and a low-pass 5th-order sinc filter, with a −3 dB attenuation at approximately 1/5th of the selected sample rate, resulting in an analog signal bandwidth of 417 Hz at the selected sampling frequency.
- The signals were acquired from 64 active electrodes mounted on an elastic cap and positioned according to the standard 10/10 coordinate system. Two additional electrodes replaced the ground electrode in conventional systems: the Common Mode Sense active electrode (CMS) and the Driven Right Leg passive electrode (DRL). The signals were recorded in single-ended mode (i.e., not referenced). Four additional electrodes were used to monitor eye movements. Specifically, two electrodes were placed on the outer canthi of both eyes to record horizontal movements (hEOG-left and hEOG-right), whereas two electrodes placed beneath and above the left eye (vEOG-lower and vEOG-upper, respectively) were used to monitor vertical movements and blinks. Finally, the signals from the left and right mastoids were also recorded to compute offline the average-mastoids reference.
- The magnitude of the offset of all electrodes was held below 20 mV.

### EEG pre-processing

- EEG signals were processed offline using custom scripts and the EEGLAB toolbox (v2022.1; [Delorme and Makeig, 2004](https://www.sciencedirect.com/science/article/abs/pii/S0165027003003479?via%3Dihub)) in the Matlab environment (R2021a, The Mathworks Inc., Natick, USA).
- Data were imported in EEGlab, re-referenced to the average of the signals recorded from left and right mastoids, resampled to 512 Hz and filtered, by applying a Hamming windowed sinc FIR filter, bandpass edges 0.5 – 40 Hz (using the function *pop_eegfiltnew* for EEGLAB, in the plugin “firfilt” v2.7.1).
- EEG signals were visually inspected to identify and remove bad channels.
- Blinks and eye-movement artifacts were corrected with a procedure based on Independent Component Analysis (ICA; see [Luck, 2022](https://socialsci.libretexts.org/Bookshelves/Psychology/Biological_Psychology/Applied_Event-Related_Potential_Data_Analysis_(Luck))). IC weights were calculated on a separate dataset, created from the original dataset by applying a bandpass filter (1-30 Hz), re-sampling to 128 Hz. The FastICA algorithm (function *fastica,* version 2.5; http://www.cis.hut.fi/projects/ica/fastica/) was used and IC weights were transferred to the original dataset.
- ICs were automatically labelled with the *pop_iclabel* function for EEGLAB, in the “ICLabel” plugin (71). Additionally, ICs accounting for blinks and/or eye movements were marked with the following procedure. (i) IC waveforms were correlated (Spearman correlation) with EOG signals. (ii) The most correlated IC with hEOG (computed as hEOG-left minus hEOG-right) was marked as IC accounting for horizontal eye movements. (iii) The most correlated IC with vEOG (computed as vEOG-upper minus vEOG-lower) was marked as IC accounting for blinks and vertical eye movements. ICs were visually inspected to confirm the selection based on the automatic labelling and the correlation with EOG channels, and removed.
- EEG epochs were extracted to compute event-related potentials (ERPs) for the encoding and the decoding epochs. Encoding epochs were extracted to assess whether action planning set a condition-specific “state” for time accumulation, before any information about probe duration is provided (e.g.: before the offset of the shortest probe). The time-locking event was the probe onset, the interval was from −200 ms to 434 ms (i.e., the shortest time interval before the probe offset) and the interval considered for baseline correction was from −200 ms to 0 ms. A voltage-based threshold of ± 100 μV was applied for bad epoch rejection. Decoding epochs were extracted to assess whether action planning affects read-out of previously encoded probe duration. The time-locking event was the probe offset, the interval was from −50 ms to 617 ms (i.e., the shortest time interval before the go signal) and the interval considered for baseline correction was from −50 ms to 50 ms (as in [Ofir and Landau, 2022](https://www.sciencedirect.com/science/article/pii/S0960982222012386?via%3Dihub)). A voltage-based threshold of ± 100 μV was applied for bad epoch rejection. We additionally monitored neural activity associated to the processing of the goal stimulus, time locked to presentation of Cued informative / Non-cued non-informative cue stimulus (.5 seconds).
- A third set of epochs was also extracted for supplementary analyses. The time-locking event was the probe onset, the interval was from −200 ms to 2000 ms (i.e., encompassing both the encoding and the decoding) and the interval considered for baseline correction was from −200 ms to 0 ms. A voltage-based threshold of ± 100 μV was applied for bad epoch rejection.

### EEG analysis

- ERPs for Encoding and Decoding were analysed separately for Direction and Distance tasks. Mass-univariate permutation-based tests, with a cluster-based method for multiple comparison correction, were implemented. Specifically, the analyses aimed at identify time points and cluster of electrodes that showed a significant effect of the Test duration (i.e., when and where ERP amplitudes differentiated among the 5 probe durations) or a significant effect of condition (i.e., when and where ERP amplitudes differentiated among Cued / Non-cued / Sensory).
- Tests were run with the function *ft_timelockstatistics* for EEGLAB, in the “Fieldtrip-lite” plugin (v20230613). Number of permutations was set to 1000, p-values for both cluster threshold and significance threshold was set to 0.01.

### Extraction of action performance features

We extracted action features (SI) from raw hand positional data in order to characterize reaching action performance and relate performance measures to timing estimates throughout trials. We parsed hand positional data to identify time at which each ballistic action started and terminated, based on a fixed velocity threshold. We defined ‘Action Latency’ as the time required to initiate each reach, therefore providing a measure of action readiness. We calculated ‘Mean Velocity’ of hand movements during hand displacement (i.e. mean velocity during interval between the start and end of each reach). We calculated ‘Reaching Error’ as the difference between the final screen position occupied by the hand at the end of each reach and the target, providing a measure of reaching precision. We extracted ‘Distance Traced’ as the product of action duration (interval between start and end of each reach) and mean velocity, indicating the amount of space covered by the reach.

### Behavioral models

#### Timing judgement and Action condition modelling

Timing perceptual judgements (proportion of “Test longer” responses) as a function of log Test durations, were fit with a Logistic function, allowing the extraction of the Point of Subjective Equality (PSE - the midpoint of the psychometric curve, identifying the Test duration that is perceptually matched to the Standard) and Just Noticeable Difference (JND - mean difference in Test duration yielding .25 and .75 proportion of “Test longer” responses, quantifying perceptual sensitivity to differences in Test durations). Linear Mixed Effects Model (LME) was used to study differences in PSE and JND, with Condition (Cued = LC, RC, LI, RI, Non-Cued and Control for Directional task; Cued = NC,FC,NI,FI, Non-Cued and Control for Distance task) as factor and participants as random effects (PSE ∼ Condition + (1 | subID); JND ∼ Condition + (1 | subID)).

#### Effects of experimental manipulations and action performance features on timing

The experimental paradigm was designed to study effects of motor preparation and plan selection on timing computations, as well as test how planned motor parameters (Directional or Distance features, congruently or incongruently mapped onto screen space) impact on timing. These questions were addressed through control of specific experimental variables. By contrast, the questions regarding the role of action performance measures on timing involve variables that were not directly controlled in the experimental design. The first concerned whether any of our performance variables (such as action latencies or the distance travelled while reaching the goal) might ultimately influence time perception. The second asks whether any such performance measure might explain the particular evolution of time responses across the duration of a block, or the entire task — reflecting an influence of learning, variations in engagement, vigour other factors on timing.

To examine the relationship between motor variables and timing judgments, we constructed a comprehensive model using the BRMS package (Bürkner, 2017). The model included the following variables: (1) probe durations, modelled non-parametrically with one free parameter per duration; (2) log-transformed action latencies; (3) a factor indicating the task being performed; (4) the line-projected distance travelled to reach the target; (5) the average movement speed throughout the reach; and (6) the precision, defined as the square root of the final distance to the target. These variables, with the addition of an intercept, interacted additively and were then logit-transformed to estimate the likelihood of the temporal judgement. The model can be run in R and is available on OSF at https://osf.io/4nz2r/.

#### Changes in time perception across trials

To assess changes in log-latencies and time responses across trials, we developed an additional Bayesian regression model using the BRMS package. This model included fixed effects for probe durations and normalized trial numbers – both within each block and across the entire cued/non-cued condition. Crucially, individual differences in the slopes of these effects were modeled as random effects. This allowed us to estimate, for each subject, their rate of change (acceleration or deceleration) in response latency and in perceived time (expansion or compression), which we could then relate statistically. For parsimony in hypothesis testing, we conflated rates of change in the Cued and Non-cued conditions (i.e. we considered these conditions as a single condition) into a single parameter (random-effect) per-subject. For log-latencies, the likelihood distribution used was Gaussian; for temporal judgements, it was a standard logistic (or inverse logit) function. The model formulas were :

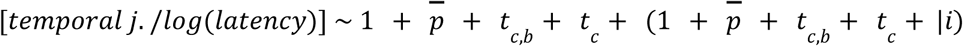

*p̄* is a vector indicating all probe durations (each modeled with its own coefficient to exclude probe-specific effects non-parametrically). *t_c,b_* is the scaled (divided by the number of last trial, e.g. 20) trial number within each block and condition. *t_c_* is the scaled (again divided by the number of last trial; Cued: 200, Non-Cued: 400) trial number within each condition. The part in parentheses signifies the individual parameter estimates which are used for post-hoc correlations, consistent with BRMS notation (*i* indicates subject number).

Given sufficient inter-individual variability, this approach enables the investigation of the relationship between the rates of change in time responses and latencies (alpha parameters that capture a linear rate of change in timing estimates and in action latency over and above the fixed effects). Spearman correlation coefficients were used to evaluate these correlations.

### Neural correlates of perceptual timing bias induced by action preparation

We identified the neural source of perceptual distortions (bias) of time occurring throughout the experiment using Representational Similarity Analysis (72) (RSA), a computational technique that allows identifying correlates of behavioural processes based on similarly patterned EEG responses. For each participant, this involved calculating timing judgement Representational Dissimilarity Matrices (Behavioural RDMs), accounting for differences (distances in Euclidean space) in proportion of “Test longer” responses across all conditions (Cued, Non-cued and Control) and test durations (Figure 5). Information structure within these matrices captures each participant’s overall performance at the timing task. This pattern reflects differences in timing judgements related to probe duration, and biases related to experimental condition. The perceptual bias in the Cued condition is reflected in asymmetries in Euclidean distances between the upper and lower triangle regions in RDM quadrants contrasting Cued against Non-cued and Sensory conditions (Figure 5a, Behavioural RDM). Neural activity was represented in an equivalent format (EEG RDMs), where for each EEG channel and 50ms time window spanning the time Encoding (440 ms following Test onset) and Decoding epochs (620 ms following Test offset) belonging to each participant, we calculated differences in neural response (Euclidean distance in average voltage) across conditions and test durations. Behavioural and EEG RDMs were compared via Kendall correlations to identify - where (which channels) and when (at what time windows) - the representational structure of neural activity resembled that of timing performance. Given that we specifically aimed at identifying the neural source of biases in perceptual time, we contrasted RDM correlations against a null model hypothesis, representing unbiased time perception. This rationale provided means of identifying channels / time windows of EEG activity that reflect biased estimates of duration, based on how they differ relative to unbiased time perception. This was achieved by generating “theoretical” timing judgement RDMs for each participant (Unbiased-Behavioural RDMs), removing timing biases within Cued, Non-cued and Control conditions. Participants’ psychometric fits were shifted so their midpoints coincided with the Standard duration (i.e. corresponding to an unbiased PSE), and proportion of “Test longer” responses were estimated across test durations based on fit parameters, thus preserving sensitivity information reflected in curve slopes. This yielded RDMs with symmetrical Euclidean distances between upper and lower triangle regions within RDM quadrants comparing different experimental conditions (Figure 5, Behavioural RDM - no-bais). Behavioural / EEG RDM r-scores and Unbiased-Behavioural / EEG RDM Kendall coefficients were then converted to z-scores (Fisher transform), and compared via t-tests (FDR corrected p-scores for multiple comparisons), yielding t-score heatmaps contrasting biased and unbiased time perception (Figure 5b,c).

## Data, Materials, and Software Availability

- All data have been uploaded to https://osf.io/4nz2r/ and are publicly available as of the date of publication.
- All original code has been uploaded to https://osf.io/4nz2r/ and is publicly available as of the date of publication.
- Any additional information required to further analyse the data reported in this paper is available from the lead contact upon request.

## Acknowledgements

This work was supported by a Marie Skłodowska-Curie Individual Fellowship (894955) awarded to NB, and an ERC Consolidator Grant (682117) awarded to DB.

## Contributions

NB conceived the study and constructed experiments. NB analyzed behavioural (timing) data, NB and MZ analyzed the EEG data, NB and FM, analyzed action data. NB, FM, MZ and DB wrote the manuscript.

## References

1. Bürkner P (2017). “brms: An R Package for Bayesian Multilevel Models Using Stan.” Journal of Statistical Software, 80(1), 1–28. doi:10.18637/jss.v080.i01

2. 10.1162/jocn_a_01459

3. Niv, Y., Daw, N., & Dayan, P. (2005). How fast to work: Response vigor, motivation and tonic dopamine. Advances in neural information processing systems, 18.

1. D. V. Buonomano, A learning rule for the emergence of stable dynamics and timing in recurrent networks. J. Neurophysiol. 94, 2275–2283 (2005).

2. J. J. Paton, D. V. Buonomano, The Neural Basis of Timing: Distributed Mechanisms for Diverse Functions. Neuron 98, 687–705 (2018).

3. R. Balasubramaniam, et al., Neural Encoding and Representation of Time for Sensorimotor Control and Learning. J. Neurosci. 41, 866–872 (2021).

4. B. Morillon, C. E. Schroeder, V. Wyart, Motor contributions to the temporal precision of auditory attention. Nat Commun 5, 5255 (2014).

5. A. Nani, et al., The Neural Correlates of Time: A Meta-analysis of Neuroimaging Studies. Journal of Cognitive Neuroscience 31, 1796–1826 (2019).

6. F. Protopapa, et al., Chronotopic maps in human supplementary motor area. PLOS Biology 17, e3000026 (2019).

7. S. Nozaradan, M. Schwartze, C. Obermeier, S. A. Kotz, Specific contributions of basal ganglia and cerebellum to the neural tracking of rhythm. Cortex 95, 156–168 (2017).

8. R. M. C. Spencer, H. N. Zelaznik, J. Diedrichsen, R. B. Ivry, Disrupted Timing of Discontinuous But Not Continuous Movements by Cerebellar Lesions. Science 300, 1437–1439 (2003).

9. H. Merchant, K. Yarrow, How the motor system both encodes and influences our sense of time. Current Opinion in Behavioral Sciences 8, 22–27 (2016).

10. A. Nuñez, W. Buño, The Theta Rhythm of the Hippocampus: From Neuronal and Circuit Mechanisms to Behavior. Frontiers in Cellular Neuroscience 15 (2021).

11. M. Iwasaki, Y. Noguchi, R. Kakigi, Neural correlates of time distortion in a preaction period. Human Brain Mapping 40, 804–817 (2019).

12. L. H. Arnal, K. B. Doelling, D. Poeppel, Delta–Beta Coupled Oscillations Underlie Temporal Prediction Accuracy. Cereb Cortex 25, 3077–3085 (2015).

13. T. W. Kononowicz, H. van Rijn, Single trial beta oscillations index time estimation. Neuropsychologia 75, 381–389 (2015).

14. M. Wiener, A. Parikh, A. Krakow, H. B. Coslett, An Intrinsic Role of Beta Oscillations in Memory for Time Estimation. Scientific Reports 8, 7992 (2018).

15. N. Hagura, R. Kanai, G. Orgs, P. Haggard, Ready steady slow: action preparation slows the subjective passage of time. Proceedings of the Royal Society B: Biological Sciences (2012).

16. M. Iwasaki, K. Tomita, Y. Noguchi, Non-uniform transformation of subjective time during action preparation. Cognition 160, 51–61 (2017).

17. I. Ayhan, D. Ozbagci, Action-induced changes in the perceived temporal features of visual events. Vision Research 175, 1–13 (2020).

18. A. Tomassini, M. Gori, G. Baud-Bovy, G. Sandini, M. C. Morrone, Motor Commands Induce Time Compression for Tactile Stimuli. J. Neurosci. 34, 9164–9172 (2014).

19. W. Zheng, H. Zhao, Y. Zhang, J. Ma, Z. Ren, Hand Movements Influence Time Perception of Visual Stimuli in Sub or Supra Seconds Duration in Engineering Psychology and Cognitive Ergonomics. Mental Workload, Human Physiology, and Human Energy, D. Harris, W.-C. Li, Eds. (Springer International Publishing, 2020), pp. 293–301.

20. P. Haggard, S. Clark, J. Kalogeras, Voluntary action and conscious awareness. Nat. Neurosci. 5, 382–385 (2002).

21. K. Yarrow, J. C. Rothwell, Manual Chronostasis: Tactile Perception Precedes Physical Contact. Current Biology 13, 1134–1139 (2003).

22. A. Tomassini, M. C. Morrone, Perceived visual time depends on motor preparation and direction of hand movements. Scientific Reports 6 (2016).

23. N. Binetti, et al., Binding space and time through action. Proceedings of the Royal Society of London B: Biological Sciences 282, 20150381 (2015).

24. R. De Kock, W. Zhou, W. M. Joiner, M. Wiener, Slowing the body slows down time perception. eLife 10, e63607 (2021).

25. G. Anobile, N. Domenici, I. Togoli, D. Burr, R. Arrighi, Distortions of visual time induced by motor adaptation. J Exp Psychol Gen 149, 1333–1343 (2020).

26. T. Yokosaka, S. Kuroki, S. Nishida, J. Watanabe, Apparent Time Interval of Visual Stimuli Is Compressed during Fast Hand Movement. PLOS ONE 10, e0124901 (2015).

27. D. Bueti, V. Walsh, The parietal cortex and the representation of time, space, number and other magnitudes. Philosophical Transactions of the Royal Society B: Biological Sciences 364, 1831–1840 (2009).

28. R. De Kock, K. A. Gladhill, M. N. Ali, W. M. Joiner, M. Wiener, How movements shape the perception of time. Trends in Cognitive Sciences 25, 950–963 (2021).

29. L. Iordanescu, M. Grabowecky, S. Suzuki, Action enhances auditory but not visual temporal sensitivity. Psychon Bull Rev 20, 108–114 (2013).

30. A. C. Fernandes, T. Garcia-Marques, The perception of time is dynamically interlocked with the facial muscle activity. Sci Rep 9, 18737 (2019).

31. D. McNamee, D. M. Wolpert, Internal Models in Biological Control. *Annual Review of Control*, Robotics, and Autonomous Systems 2, 339–364 (2019).

32. D. W. Franklin, D. M. Wolpert, Computational Mechanisms of Sensorimotor Control. Neuron 72, 425–442 (2011).

33. M. Joch, M. Hegele, H. Maurer, H. Müller, L. K. Maurer, Accuracy of Motor Error Predictions for Different Sensory Signals. Front. Psychol. 9 (2018).

34. J. W. Krakauer, M.-F. Ghilardi, C. Ghez, Independent learning of internal models for kinematic and dynamic control of reaching. Nat Neurosci 2, 1026–1031 (1999).

35. G. Gavazzi, A. Bisio, T. Pozzo, Time perception of visual motion is tuned by the motor representation of human actions. Scientific reports 3 (2013).

36. R. De Kock, W. Zhou, P. Datta, W. Mychal Joiner, M. Wiener, The role of consciously timed movements in shaping and improving auditory timing. Proceedings of the Royal Society B: Biological Sciences 290, 20222060 (2023).

37. K. Yarrow, P. Haggard, R. Heal, P. Brown, J. C. Rothwell, Illusory perceptions of space and time preserve cross-saccadic perceptual continuity. Nature 414, 302–305 (2001).

38. Z. Zhang, D. Sternad, The primacy of rhythm: how discrete actions merge into a stable rhythmic pattern. Journal of Neurophysiology 121, 574–587 (2019).

39. J. Guo, Z. Zhang, D. Sternad, J.-H. Song, Improved motor timing enhances time perception. Journal of Vision 19, 218b (2019).

40. H. Merchant, W. Zarco, L. Prado, Do we have a common mechanism for measuring time in the hundreds of millisecond range? Evidence from multiple-interval timing tasks. J. Neurophysiol. 99, 939 (2008).

41. H. Merchant, O. Pérez, W. Zarco, J. Gámez, Interval tuning in the primate medial premotor cortex as a general timing mechanism. The Journal of Neuroscience 33, 9082–9096 (2013).

42. N. Binetti, A. Tomassini, K. Friston, S. Bestmann, Uncoupling Sensation and Perception in Human Time Processing. Journal of Cognitive Neuroscience 32, 1369–1380 (2020).

43. M. C. Morrone, J. Ross, D. Burr, Saccadic eye movements cause compression of time as well as space. Nat. Neurosci. 8, 950–954 (2005).

44. A. G. Chowdhary, J. H. Challis, Timing Accuracy in Human Throwing. Journal of Theoretical Biology 201, 219–229 (1999).

45. M. Wittmann, V. Van Wassenhove, B. Craig, M. P. Paulus, The neural substrates of subjective time dilation. Front. Hum. Neurosci. 4 (2010).

46. K. E. Adolph, J. E. Hoch, Motor Development: Embodied, Embedded, Enculturated, and Enabling. Annual Review of Psychology 70, 141–164 (2019).

47. A. Benedetto, P. Binda, M. Costagli, M. Tosetti, M. C. Morrone, Predictive visuo-motor communication through neural oscillations. Current Biology 31, 3401–3408.e4 (2021).

48. A. Tomassini, L. Ambrogioni, W. P. Medendorp, E. Maris, Theta oscillations locked to intended actions rhythmically modulate perception. eLife 6, e25618 (2017).

49. S.-J. Blakemore, D. M. Wolpert, C. D. Frith, Central cancellation of self-produced tickle sensation. Nat Neurosci 1, 635–640 (1998).

50. P. Cisek, Making decisions through a distributed consensus. Current Opinion in Neurobiology 22, 927–936 (2012).

51. L. Shmuelof, J. W. Krakauer, P. Mazzoni, How is a motor skill learned? Change and invariance at the levels of task success and trajectory control. Journal of Neurophysiology 108, 578–594 (2012).

52. G. Scozia, et al., Space is a late heuristic of elapsing time: New evidence from the STEARC effect. Cortex 164, 21–32 (2023).

53. G. Scozia, et al., The time course of the spatial representation of ‘past’ and ‘future’ concepts: New evidence from the STEARC effect. Atten Percept Psychophys 86, 1048–1055 (2024).

54. A. Tomassini, D. Spinelli, M. Jacono, G. Sandini, M. C. Morrone, Rhythmic Oscillations of Visual Contrast Sensitivity Synchronized with Action. J. Neurosci. 35, 7019–7029 (2015).

55. J. Diedrichsen, K. Kornysheva, Motor skill learning between selection and execution. Trends in Cognitive Sciences 19, 227–233 (2015).

56. H. Pashler, Shifting visual attention and selecting motor responses: Distinct attentional mechanisms. Journal of Experimental Psychology: Human Perception and Performance 17, 1023–1040 (1991).

57. F. Moscatelli, et al., Relationship between Cortical Excitability and Complex Reaction Time. THE OPEN NEUROLOGY JOURNAL 17 (2023).

58. P. Mazzoni, A. Hristova, J. W. Krakauer, Why Don’t We Move Faster? Parkinson’s Disease, Movement Vigor, and Implicit Motivation. J. Neurosci. 27, 7105–7116 (2007).

59. Y. Niv, N. Daw, P. Dayan, How fast to work: Response vigor, motivation and tonic dopamine in Advances in Neural Information Processing Systems, (MIT Press, 2005).

60. P. Dayan, Instrumental vigour in punishment and reward. European Journal of Neuroscience 35, 1152–1168 (2012).

61. A. Betancourt, O. Pérez, J. Gámez, G. Mendoza, H. Merchant, Amodal population clock in the primate medial premotor system for rhythmic tapping. Cell Reports 42, 113234 (2023).

62. J. Wang, D. Narain, E. A. Hosseini, M. Jazayeri, Flexible timing by temporal scaling of cortical responses. Nat Neurosci 21, 102–110 (2018).

63. S. Zhou, S. C. Masmanidis, D. V. Buonomano, Neural Sequences as an Optimal Dynamical Regime for the Readout of Time. Neuron 108, 651–658.e5 (2020).

64. P. Salvioni, M. M. Murray, L. Kalmbach, D. Bueti, How the visual brain encodes and keeps track of time. J. Neurosci. 33, 12423–12429 (2013).

65. V. Centanino, G. Fortunato, D. Bueti, The neural link between stimulus duration and spatial location in the human visual hierarchy. Nat Commun 15, 10720 (2024).

66. V. Pouthas, et al., Neural network involved in time perception: an fMRI study comparing long and short interval estimation. Hum. Brain Mapp. 25, 433–441 (2005).

67. D. Bueti, E. Macaluso, Physiological correlates of subjective time: evidence for the temporal accumulator hypothesis. Neuroimage 57, 1251–1263 (2011).

68. A. Damsma, N. Schlichting, H. van Rijn, Temporal Context Actively Shapes EEG Signatures of Time Perception. J. Neurosci. 41, 4514–4523 (2021).

69. N. Ofir, A. N. Landau, Neural signatures of evidence accumulation in temporal decisions. Current Biology 32, 4093–4100 (2022).

70. A. Tomassini, T. Vercillo, F. Torricelli, M. C. Morrone, Rhythmic motor behaviour influences perception of visual time. Proceedings of the Royal Society B: Biological Sciences 285, 20181597 (2018).

71. L. Pion-Tonachini, K. Kreutz-Delgado, S. Makeig, ICLabel: An automated electroencephalographic independent component classifier, dataset, and website. NeuroImage 198, 181–197 (2019).

72. N. Kriegeskorte, M. Mur, P. A. Bandettini, Representational similarity analysis - connecting the branches of systems neuroscience. Front. Syst. Neurosci. 2 (2008).

