## Supplemental Figures for "Evolution of visual time throughout goal driven action learning"

* dual senior authorship

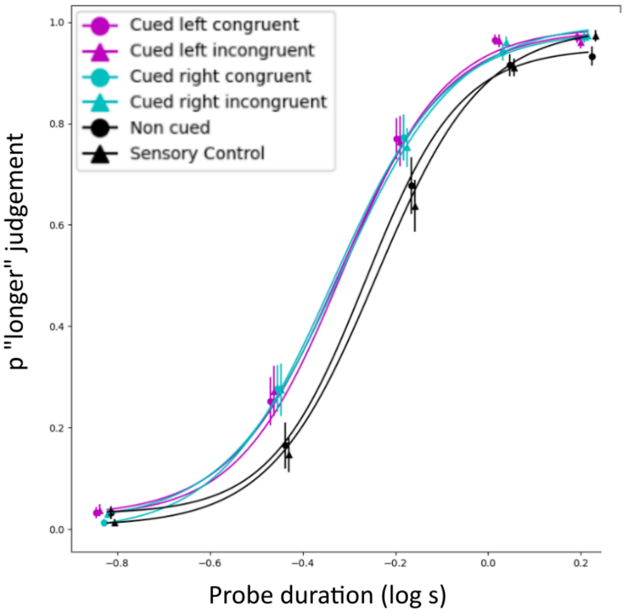


Directional task contrasts:

| contrast | estimate | SE | df | t.ratio | p.value |
|---|---|---|---|---|---|
| LC - LI | -0.001875 | 0.0133 | 95 | -0.141 | 1.0000 |
| LC - NonCued | -0.038807 | 0.0133 | 95 | -2.910 | 0.0498 |
| LC - RC | 0.000199 | 0.0133 | 95 | 0.015 | 1.0000 |
| LC - RI | 0.003784 | 0.0133 | 95 | 0.284 | 0.9997 |
| LC - Sensory | -0.053774 | 0.0133 | 95 | -4.033 | 0.0015 |
| LI - NonCued | -0.036932 | 0.0133 | 95 | -2.770 | 0.0713 |
| LI - RC | 0.002074 | 0.0133 | 95 | 0.156 | 1.0000 |
| LI - RI | 0.005659 | 0.0133 | 95 | 0.424 | 0.9982 |
| LI - Sensory | -0.051899 | 0.0133 | 95 | -3.892 | 0.0025 |
| NonCued - RC | 0.039006 | 0.0133 | 95 | 2.925 | 0.0478 |
| NonCued - RI | 0.042591 | 0.0133 | 95 | 3.194 | 0.0227 |
| NonCued - Sensory | -0.014967 | 0.0133 | 95 | -1.123 | 0.8709 |
| RC - RI | 0.003585 | 0.0133 | 95 | 0.269 | 0.9998 |
| RC - Sensory | -0.053973 | 0.0133 | 95 | -4.048 | 0.0014 |
| RI - Sensory | -0.057558 | 0.0133 | 95 | -4.317 | 0.0005 |

**Figure S1**: Psychometric fit and PSE (point of subjective equality) LME ANOVA contrasts of 6 conditions (4 Cued, Non-cued and Sensory control) - Direction task. Cued trials (target is shown during foreperiod) are split in 4 types based on action type (left or right hand reach) and hand to screen space mapping (congruent or incongruent): LC = Left Congruent; RC = Right Congruent; LI = Left Incongruent (right hand movement to reach Left target); RI = Reft Incongruent (left hand movement to reach Right target). NonCued (target is not shown during foreperiod, and its presentation coincides with the Go signal). Sensory (Test is timed in matched interval where action is not prepared).

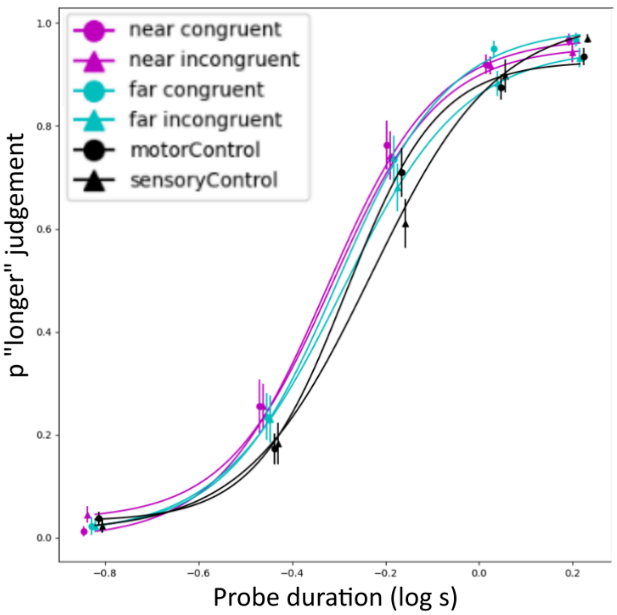


Distance task contrasts:

| contrast | estimate | SE | df | t.ratio | p.value |
|---|---|---|---|---|---|
| FC - FI | -0.00788 | 0.0173 | 95 | -0.457 | 0.9974 |
| FC - NC | -0.00990 | 0.0173 | 95 | -0.574 | 0.9925 |
| FC - NI | -0.00419 | 0.0173 | 95 | -0.243 | 0.9999 |
| FC - NonCued | -0.02829 | 0.0173 | 95 | -1.639 | 0.5748 |
| FC - Sensory | -0.05745 | 0.0173 | 95 | -3.330 | 0.0152 |
| FI - NC | -0.00202 | 0.0173 | 95 | -0.117 | 1.0000 |
| FI - NI | 0.00369 | 0.0173 | 95 | 0.214 | 0.9999 |
| FI - NonCued | -0.02041 | 0.0173 | 95 | -1.183 | 0.8440 |
| FI - Sensory | -0.04958 | 0.0173 | 95 | -2.873 | 0.0548 |
| NC - NI | 0.00570 | 0.0173 | 95 | 0.331 | 0.9995 |
| NC - NonCued | -0.01839 | 0.0173 | 95 | -1.066 | 0.8936 |
| NC - Sensory | -0.04756 | 0.0173 | 95 | -2.756 | 0.0738 |
| NI - NonCued | -0.02409 | 0.0173 | 95 | -1.396 | 0.7291 |
| NI - Sensory | -0.05326 | 0.0173 | 95 | -3.087 | 0.0308 |
| NonCued - Sensory | -0.02917 | 0.0173 | 95 | -1.691 | 0.5414 |

**Figure S2**: Psychometric fit and PSE (point of subjective equality) LME ANOVA contrasts of 6 conditions (4 Cued, Non-cued and Sensory control) - Distance task. Cued trials (target is shown during foreperiod) are split in 4 types based on action type (rightward near or far hand reach) and hand to screen space mapping (congruent or incongruent): NC = Near Congruent; FC = Far Congruent; NI = Near Incongruent (large hand movement to reach Near target); FI = Far Incongruent (small movement to reach Far target).

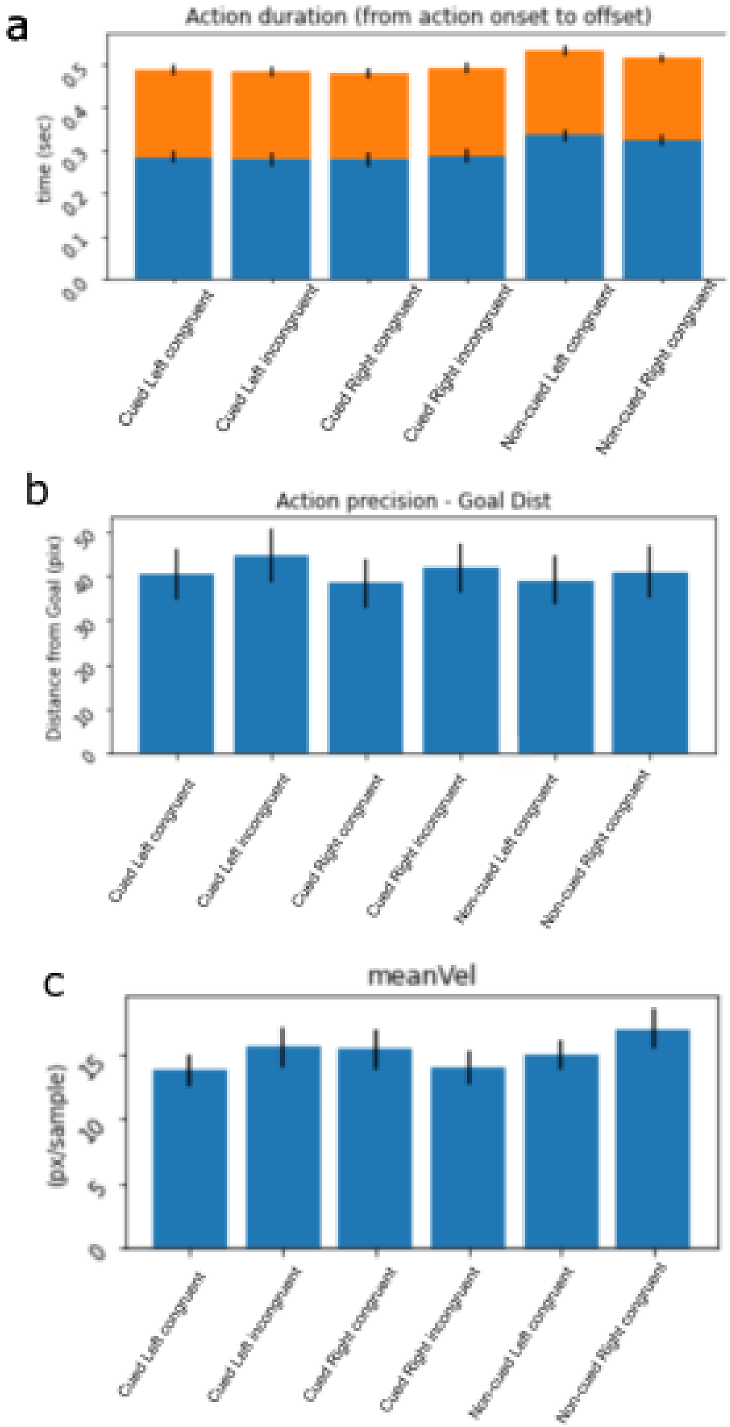


**Figure S3:** Direction task action features across 6 experimental conditions. a) Action latency (blue) and duration (orange), respectively measured as interval between go signal and action start, and interval between action start and action end. b)  Reaching errors across conditions, measured as distance between landing point and target center.  c) Mean action velocity.

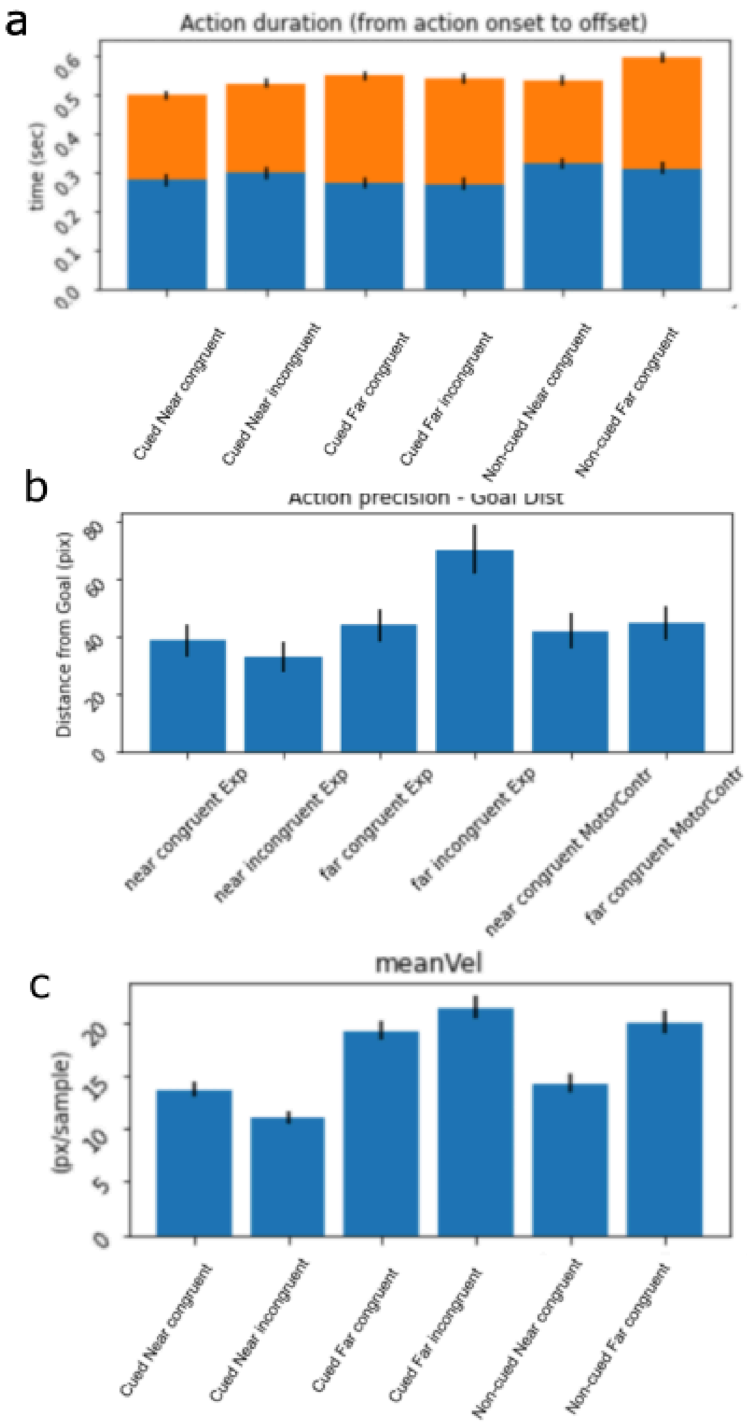


**Figure S4:** Distance task action features across 6 experimental conditions. a) Action latency (blue) and duration (orange). b) Reaching errors across conditions, measured as distance between landing point and target center. c) Mean action velocity.

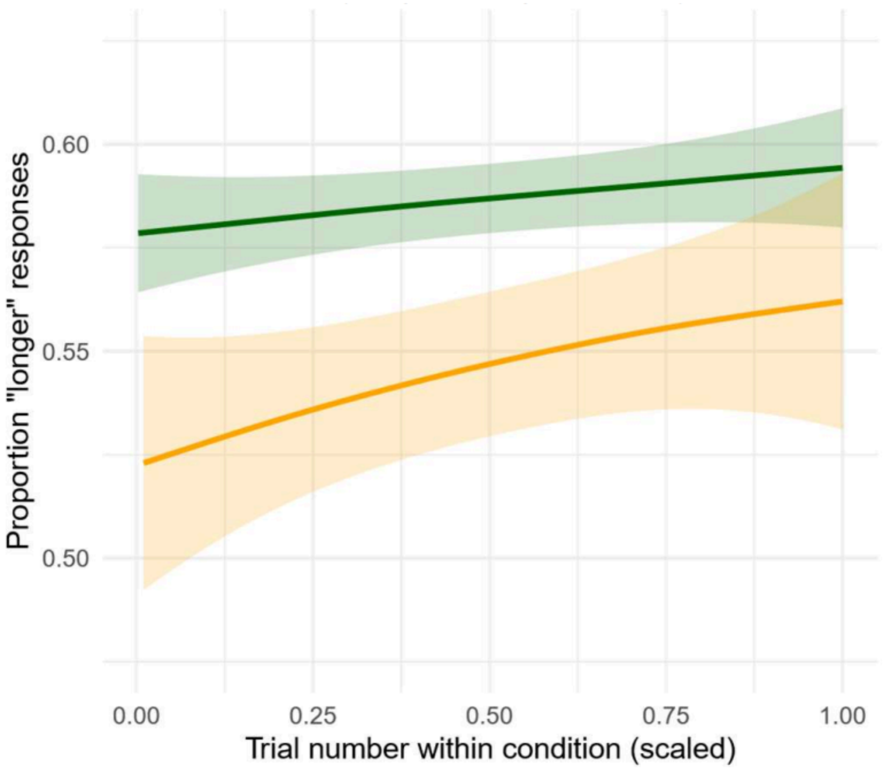


**Figure S5**: Timing judgements as a function of trial number, showing progressive expansion of perceived time. Rates of change in timing judgements were individually modelled for the Cued (green) and Non-cued (orange) conditions. The differences in intercept between these curves reflect the overall difference in PSEs estimating through psychometric fits.

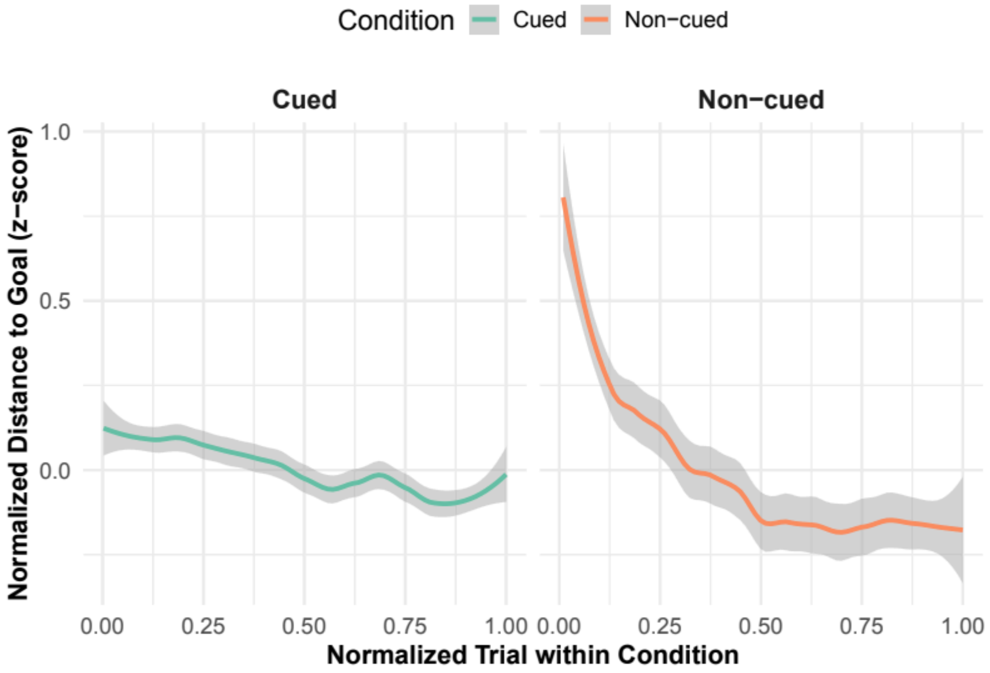


**Figure S6**: Hand reach precision (normalized distance from goal area centre) as a function of trial number, showing progressive improvement in action precision. Rates of change in timing judgements were individually modelled for the Cued (green) and Non-cued (orange) conditions. Rates of change in precision are more pronounced in Non-cued trials, reflecting additional cost in handling target location uncertainty.

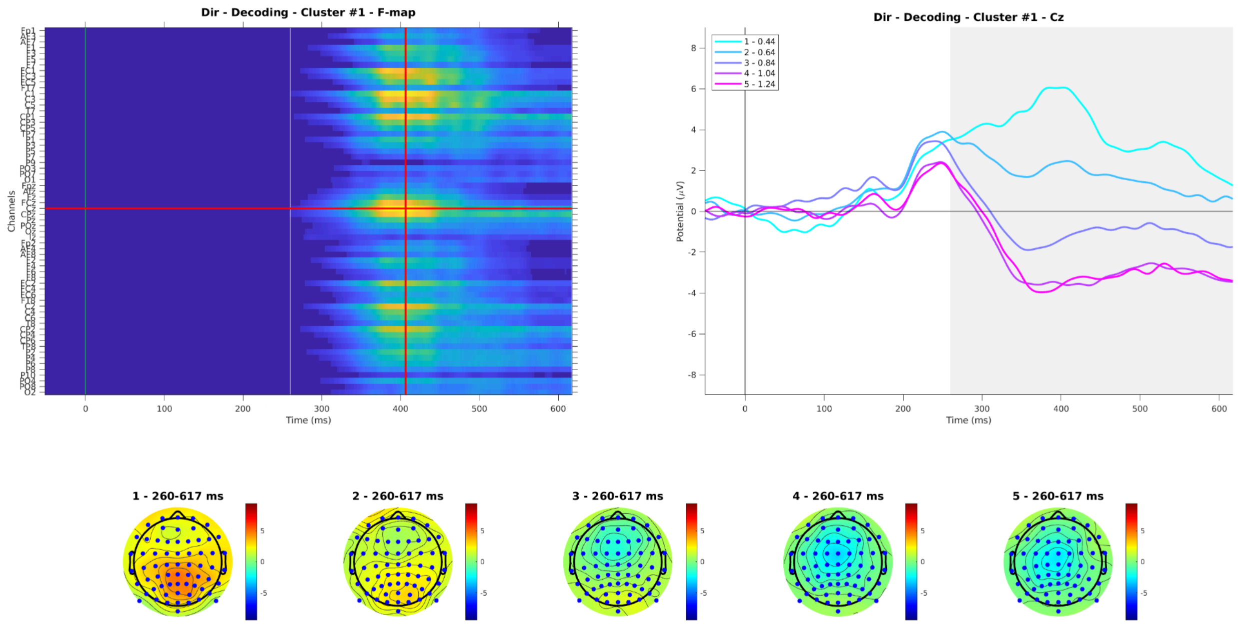


**Figure S7**: Direction Task Time Decoding ERP analysis - Effect of Probe at electrode Cz

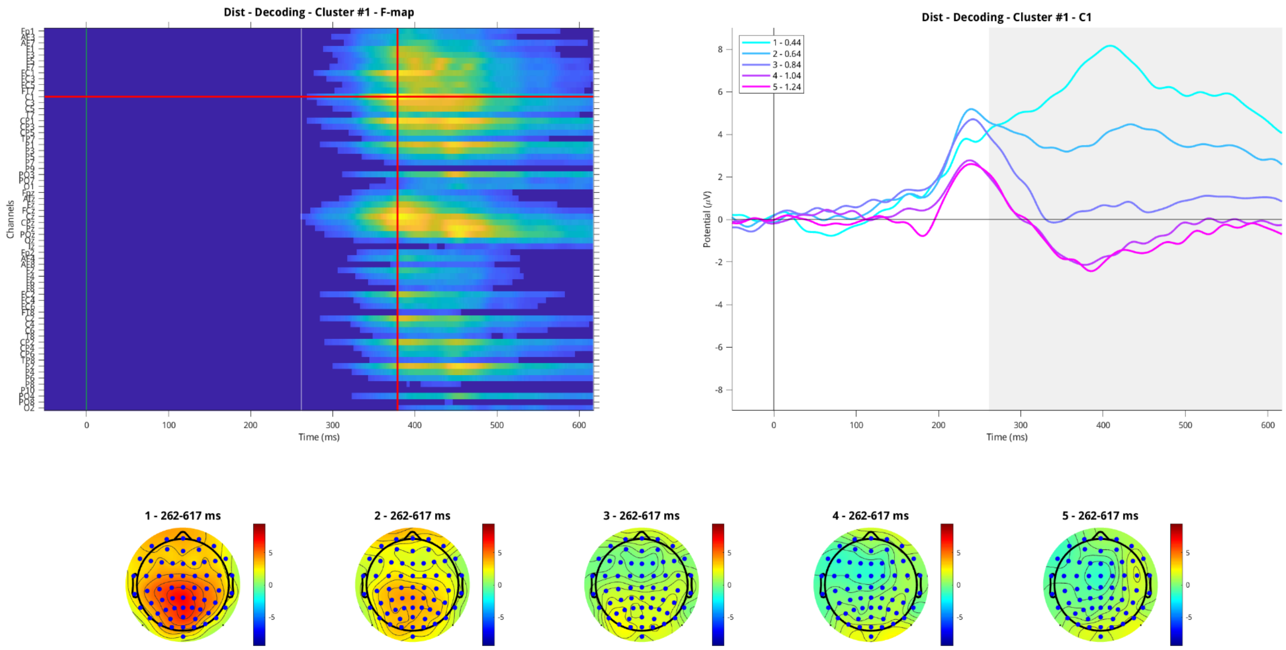


**Figure S8**: Distance Task Time Decoding ERP analysis - Effect of Probe at electrode C1

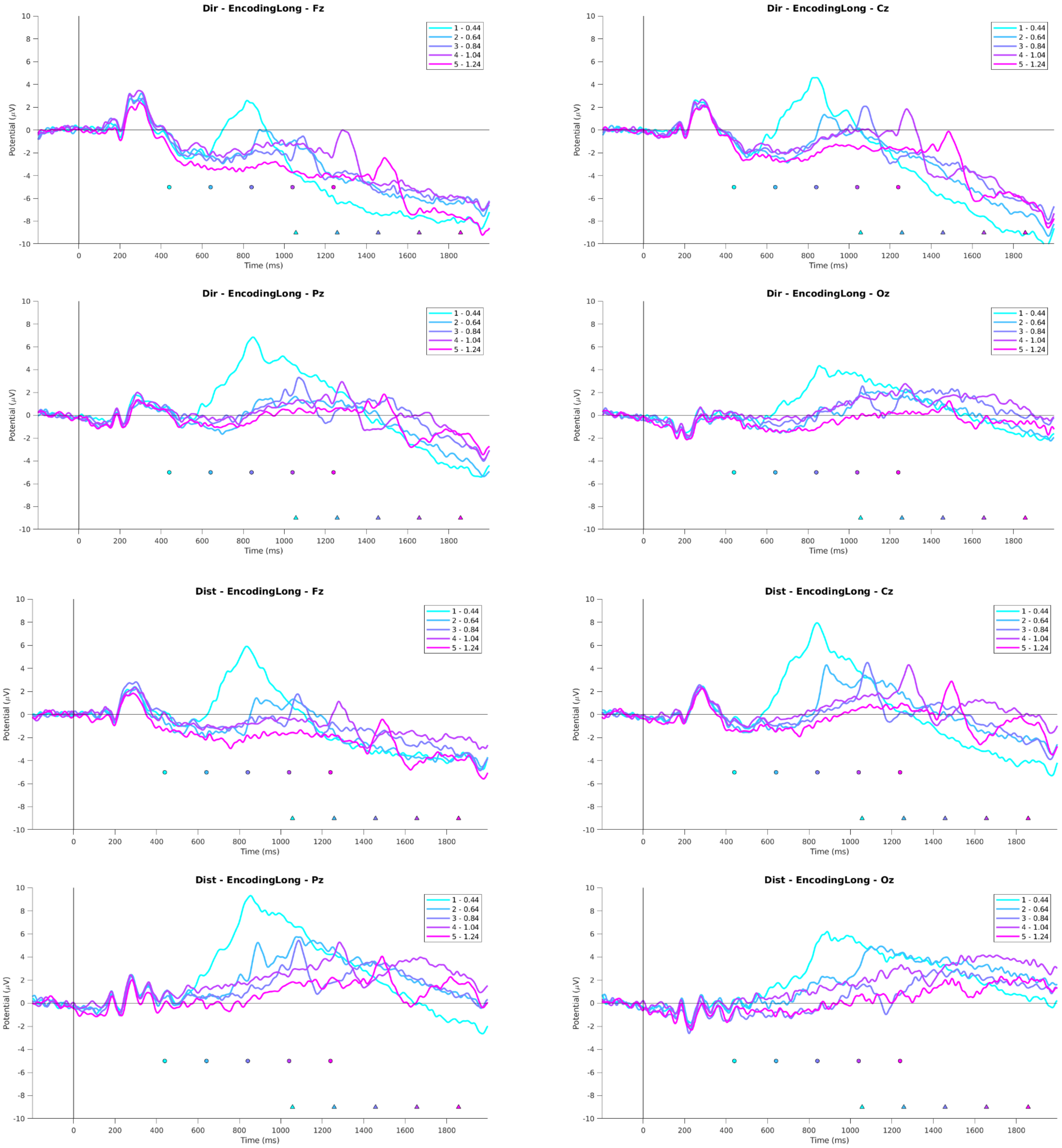


**Figure S9:** ERP epochs encompassing time encoding and decoding. Direction task in top two rows, Distance task in bottom two rows. Coloured dots represent probe offsets. To note, in the Decoding epochs, these time points represent the 0-ms point and the interval for baseline correction ranges from 50 ms before to 50 ms after probe offset. Coloured triangles represent the shortest presentation of the GO signal, relative to the corresponding probe offset.

***Cued Vs Non-cued***

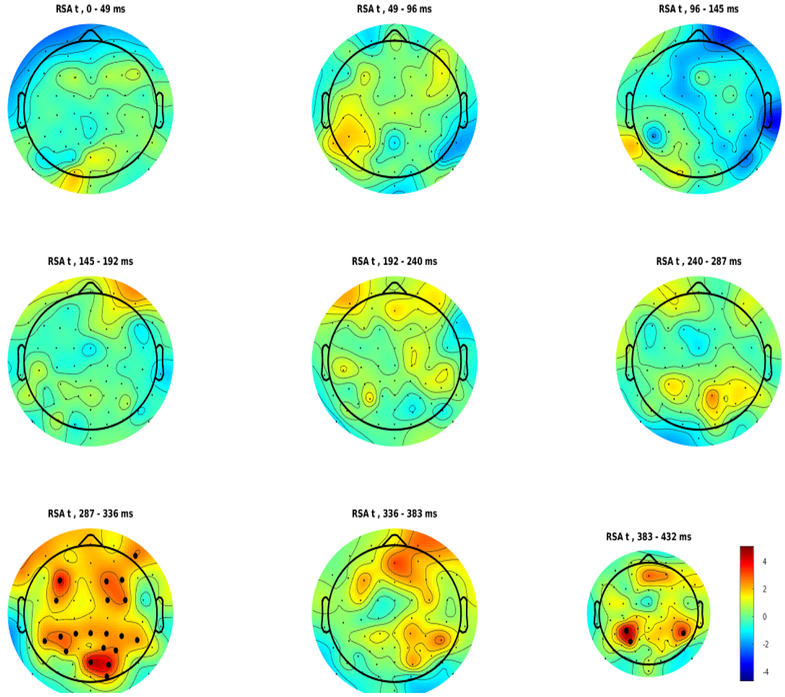


***Cued Vs Sensory***

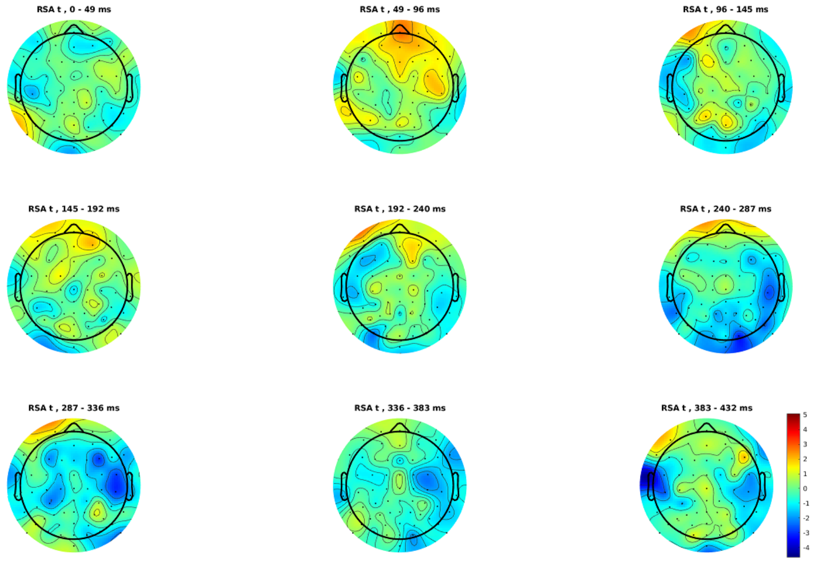


**Figure S10:** Direction Task - Encoding epoch RSA analysis (.43 sec following test stimulus onset), comparing Cued relative to Non-cued and Sensory conditions.

#### *Cued Vs Non-cued*

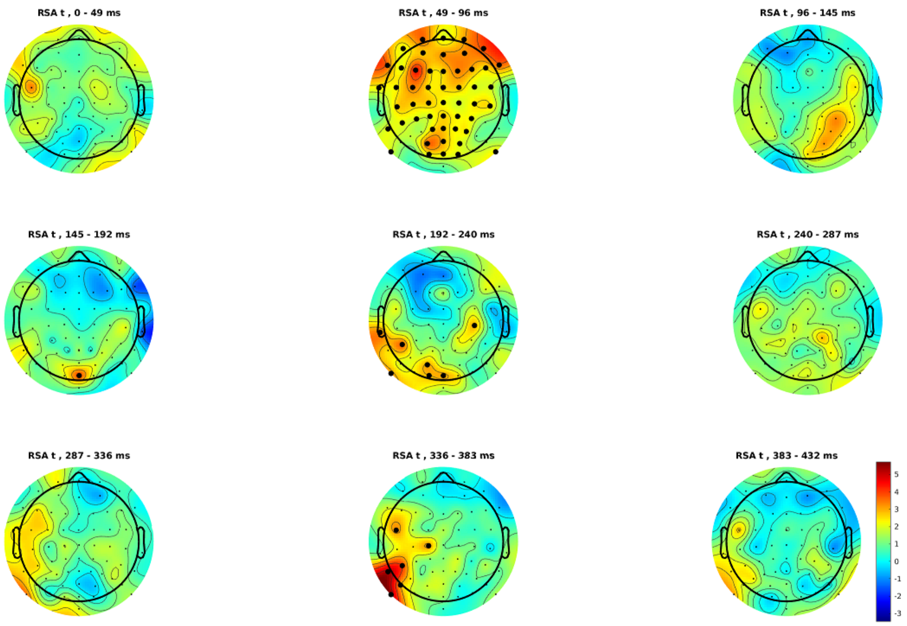


#### *Cued Vs Sensory*

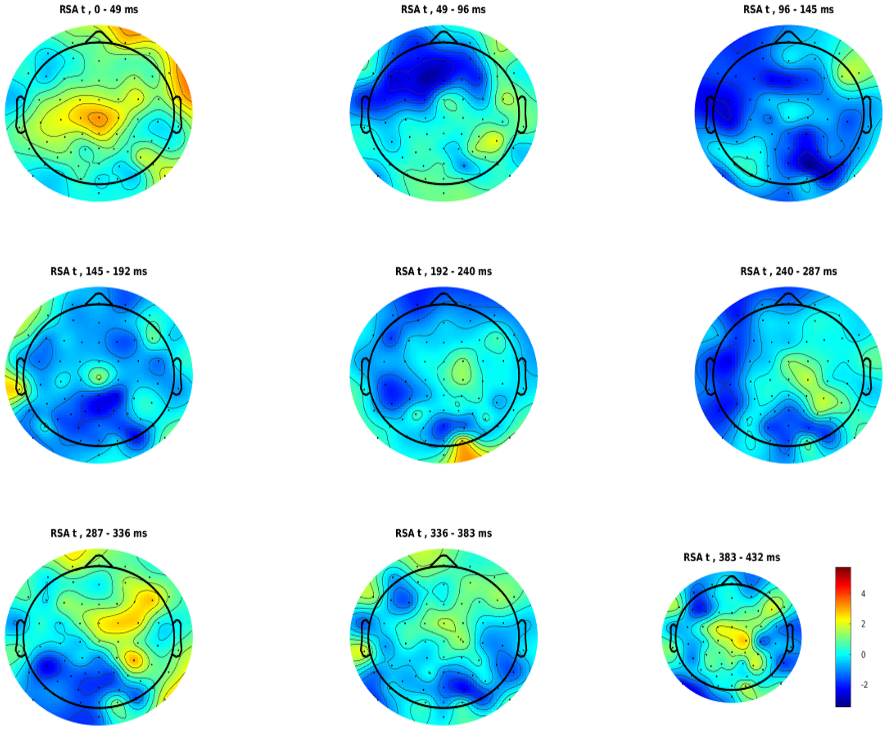


**Figure S11:** Distance Task - Encoding epoch RSA analysis (.43 sec following test stimulus onset), comparing Cued relative to Non-cued and Sensory conditions.

### *Cued Vs Non-cued*

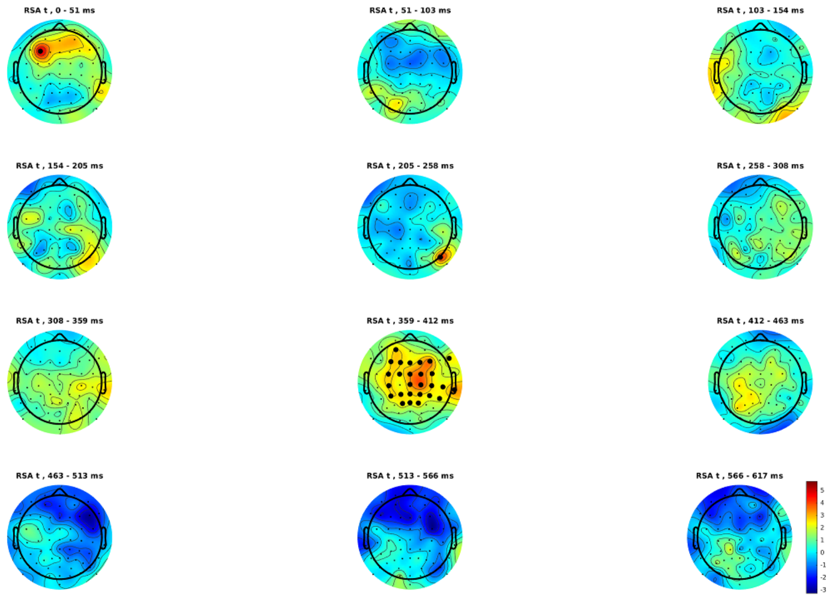


### *Cued Vs Sensory*

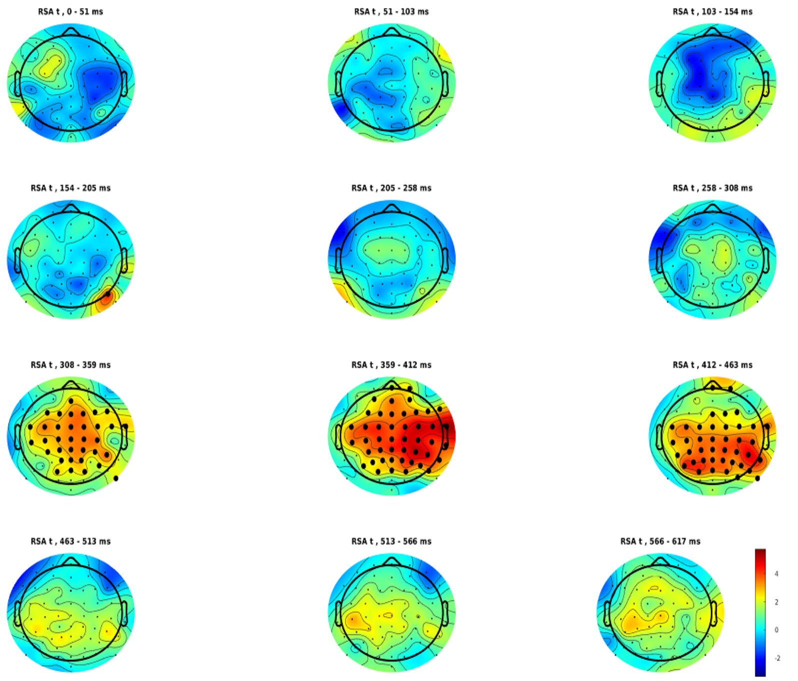


**Figure S12:** Direction Task - Decoding epoch RSA analysis (.62 sec following test stimulus onset), comparing Cued relative to Non-cued and Sensory conditions.

***Cued Vs Non-cued***

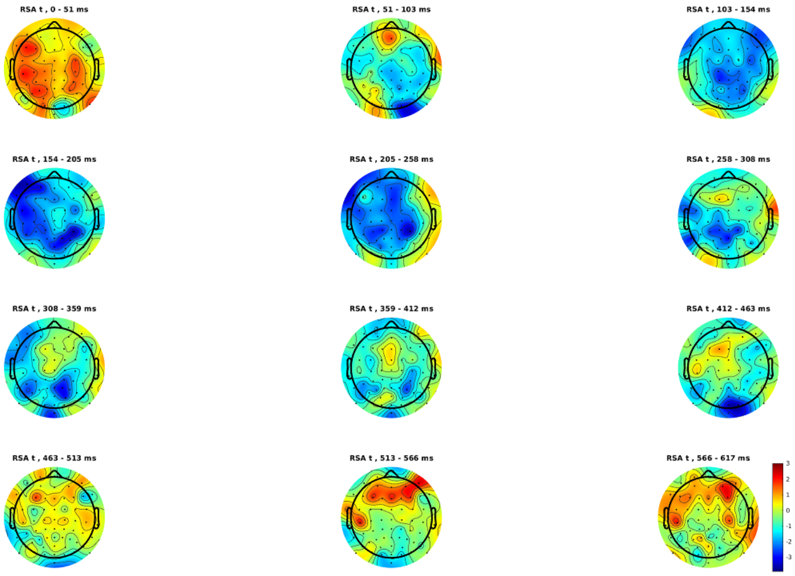


***Cued Vs Sensory***

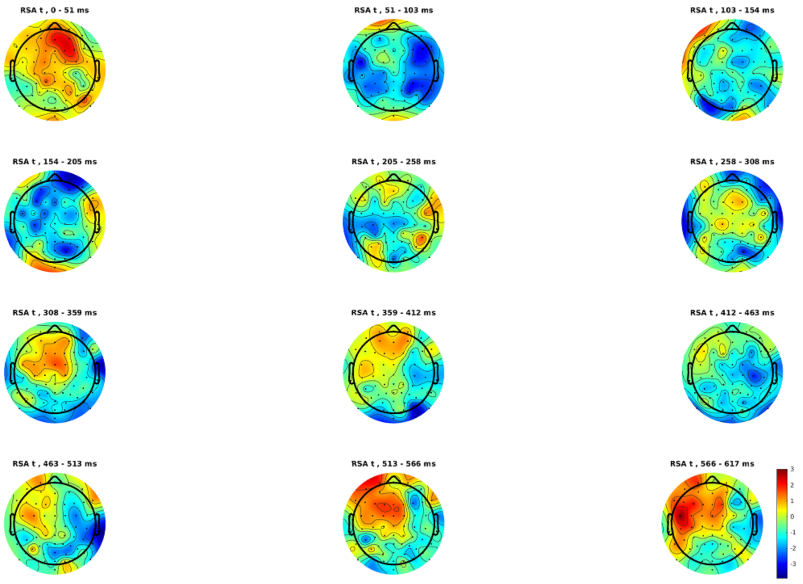


**Figure S13:** Distance Task - Decoding epoch RSA analysis (.62 sec following test stimulus onset), comparing Cued relative to Non-cued and Sensory conditions.
